# Cell-state transitions and frequency-dependent interactions among subpopulations together explain the dynamics of spontaneous epithelial-mesenchymal heterogeneity in breast cancer

**DOI:** 10.1101/2023.12.07.567986

**Authors:** Paras Jain, Ramanarayanan Kizhuttil, Madhav B Nair, Sugandha Bhatia, Erik W Thompson, Jason T. George, Mohit Kumar Jolly

**Author notes:** Authors to whom correspondence should be addressed (J.T.G.), (M.K.J.). These authors contributed equally to the study (Paras Jain, Ramanarayanan Kizhuttil).

## Abstract

Individual cells in a tumour can be distributed among Epithelial (E) and Mesenchymal (M) cell-states, as characterised by the levels of canonical E and M markers. Even after E and M (E-M) subpopulations are isolated and then cultured independently, E-M heterogeneity can re-equilibrate in each population over time, sometimes regaining the initial distribution of the parental cell population. However, it remains unclear which population-level processes give rise to the dynamical changes in E-M heterogeneity observed experimentally, including 1) differential growth, 2) cell-state switching, and 3) frequency-dependent growth or state-transition rates. Here, we analyse the necessity of these three processes in explaining the dynamics of E-M population distributions as observed in PMC42-LA and HCC38 breast cancer cells. We find that growth differences among E and M subpopulations, with and without any frequency-dependent interactions (cooperation or suppression) among E-M sub-populations, are insufficient to explain the observed population dynamics. This insufficiency is ameliorated by including cell-state transitions, albeit at slow rates, in explaining both PMC42-LA and HCC38 cells data. Further, our models predict that treatment of HCC38 cells with TGFβ signalling and JAK2/3 inhibitors could significantly enhance the transition rates from M state to E state, but does not prevent transitions from E to M. Finally, we devise a selection criterion to identify the next most informative time points for which future experimental data can optimally improve the identifiability of our estimated best fit model parameters. Overall, our study identifies the necessary population-level processes shaping the dynamics of E-M heterogeneity in breast cancer cells.

## Introduction

Tumour heterogeneity still remains a major challenge in treating cancer. Population-level heterogeneity manifests as the occurrence of distinct cellular phenotypes resulting from genetic and non-genetic variability amongst single cells. While extensive genome-level characterisation of many cancer types has identified driver mutations and directed the development of targeted drugs to contain tumour size, the emergence of resistance and tumour relapse is an ever-present threat. Drug-resistant populations emerge under evolutionary pressures that ultimately select for slowly proliferating pre-existent resistant cells or acquired resistance^1^. Recent empirical evidence has suggested both pre-existent and acquired resistance can be found even in clonal populations with cells having identical genetic backgrounds ^2^. Non-genetic heterogeneity and its consequent resistance therefore further complicates our understanding given the diversity of levels (proteome, metabolome) and cellular processes (Epithelial-Mesenchymal transition, stem cell/non-stem cell, glycolysis/oxidative phosphorylation) at which cells can manifest epigenetic (non-genetic) differences^3^.

Switching among E and M states represents one canonical example of non-genetic cellular heterogeneity and has been reported in embryonic development, wound healing, and diseased states, including fibrosis and cancer metastasis. The switches among E-M states have been shown to occur by mutual interaction between several regulatory players and signalling pathways ^4,5^. Further, single-cell characterisation at the transcriptome and chromatin levels has revealed the extent of E-M transitions (EMT) and M-E transitions (MET) in response to different concentrations of growth factors and the duration of their exposure ^6–8^. Apart from molecular characterisation of EMT and MET, few population-level studies exist to evaluate E-M state composition of cancer cells and their corresponding changes over time. The E and M cells are found to be distributed in variable fractions across distinct cell lines, even within the same cancer type. For example, luminal-like and claudin-low breast cancer cell lines were shown to be comprised of a majority of epithelial and mesenchymal cells, while basal-like cell lines were enriched in cells co-expressing E and M markers ^9^. The phenotypic composition of E-M cells can also change in experiments tracking clonal populations of single cells derived from a parental cell line ^10^. Similarly, the PMC42-LA breast cancer cell line exhibits a stable composition (80:20 ratio of E:M cells), however its parental cell line PMC42-ET is M cell-dominant ^11,12^. Consequently, this variability necessitates a more detailed framework capable of explaining such differences, yet the mechanisms determining phenotypic composition at a population level and its dynamical evolution currently remain unknown.

Experimental evidence suggests temporally varying E-M cell fractions when E-M populations are either isolated or mixed in different proportions ^9,12–14^. E, M, and hybrid E/M subpopulations (using canonical E-Cadherin and Vimentin levels) were simultaneously observed in castration-resistant metastatic prostate cancer cell lines ^13^. Inhibition of HMGA2-destabilized M phenotypic state and reduced E-M transition thereby lowering down the proportion of M cells in the population. Similarly, in HCC38 breast cancer cells, the phenotypic composition of E-M cells in the population was found to influence the extent of EMT and/or MET. Further, the inhibition of the TGFβ signalling pathway and blocking of JAK2/3 diminished the effects that the M cells had on EMT. Thus, the authors suggested cell-cell communication was at least partly responsible for influencing cell-state transitions at the population level. Unfortunately, however, the above studies did not explicitly report the growth rates of E-M cell populations in each scenario, thus missing out inadvertently on contributions of different growth rates on E-M heterogeneity dynamics. Measuring any change(s) in growth rates is crucial for two reasons: 1) growth rate differences among subpopulations impact the population-level composition ^15,16^, and 2) EMT is also known to slows down cell proliferation ^17,18^, and thus, inhibition of EMT may not only reduce cell-state transition but may increase the proliferation rate^19^. Therefore, the relative contribution of various interconnected cell-autonomous and non-cell autonomous effects on population dynamics of EMT and MET remains poorly understood.

Here, we investigated which population-level processes are necessary to explain the experimentally observed time-course data on spontaneous emergence of E-M heterogeneity in PMC42-LA and HCC38 breast cancer lines - 1) differential growth rates, 2) cell-state transitions, and 3) cell-cell communication via modulating either growth or cell-state transition rates ^9,12^. We compared a number of distinct mathematical models combining one or more of the above cellular processes according to their goodness of fit to the experimental data, while optimizing model parameterization restricted to biologically relevant ranges. Applying the information theoretic-based Corrected Akaike Information Criteria (AICc) to select the best fit model, we found that including E-M cell state transitions was necessary, albeit at slow rates, to explain the emergence of dynamic E-M heterogeneity observed in both PMC42-LA and HCC38 cells. Further, on incorporating cell-state transitions, the models having frequency-dependent growth rates were better in explaining the data than the models with frequency-dependent cell-state transition rates. The diminished role of cellular interactions influencing cell-state transitions was substantiated by fitting TGFβ- and JAK2/3-inhibitor treatment data as well. We performed uncertainty analysis of models in explaining the data by commenting on their false positive selection due to noisy experimental data and sensitivities to the initial proportion of E and M cells in the experiments. Lastly, we devised a selection criterion to identify the next most informative time points for which future experimental data optimally improve the identifiability (95% confidence bounds) of our estimated best fit model parameters. Overall, our results demonstrate how phenotypic heterogeneity in breast cancer cells emerges as a consequence of cell state transitions together with the influence of heterogeneous subpopulations on each subpopulation’s growth rate.

## Results

### Cell-state transitions are necessary for explaining spontaneous EMP in PMC42-LA breast cancer cells

Recent experiments reported that PMC42-LA breast cancer cells exhibited an 80:20 ratio of E and M subpopulations based on Epithelial Cell Adhesion Molecule (EpCAM) levels (EpCAM^high^ and EpCAM^low^, respectively) ^12^. When both subpopulations were isolated and cultured independently, they recapitulated the original parental distribution of an 80:20 E:M ratio after eight weeks of time (culture 1 and 2 in **Figure 1A** – Exp data). These observations of spontaneous E-M plasticity (EMP) motivated us to investigate the role of distinct population-level processes (differential growth, cell-state transition, cell-cell communication) in governing these dynamics. Thus, we designed a cohort of mathematical models, each including different combinations of these processes and compared their fit with the experimental data.

**Figure 1.**
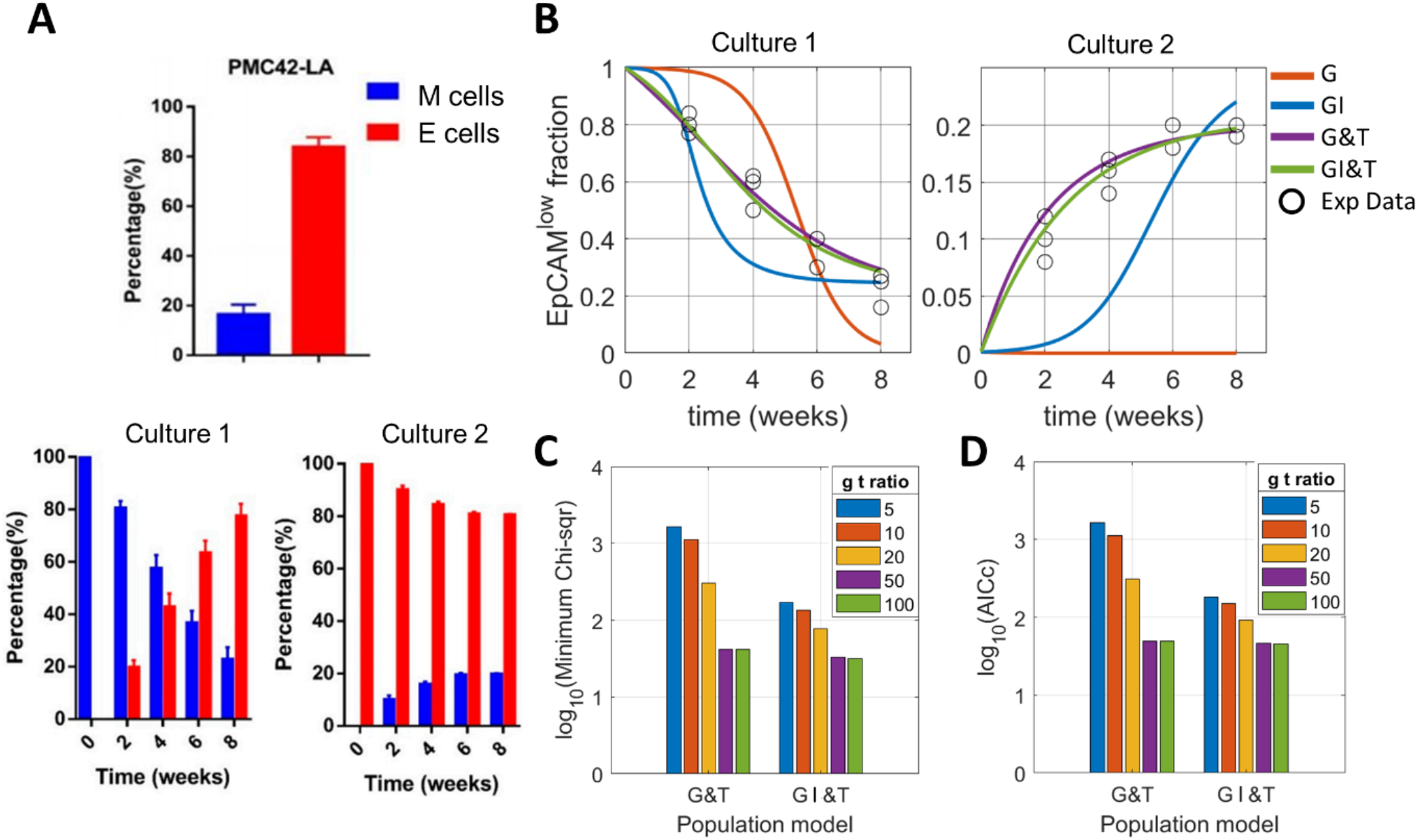
Necessity of cell-state transition in explaining spontaneous EMP in PMC42-LA breast cancer cells. **A)** Distribution of EpCAM^high^ (epithelial) ^and^ EpCAM^low^ (mesenchymal) cells in PMC42-LA cell; **ii)** changing phenotypic distribution of the population in cell cultures of isolated E and M cells from the parental cell line. 100% M and 100% E cells at week 0 in culture 1 and 2, respectively (Figure adopted from Bhatia et al. 2019 ^12^). **B)** Fits to the experimental data using population models that consider – 1) Growth of E-M subpopulations (G – growth), 2) E-M subpopulations mutual influence on each other’s growth (GI – growth influence), 3) Growth and cell-state transition among E-M subpopulations (G&T – growth and transition), and 4) E-M subpopulations influence on each other’s growth with cell-state transitions (GI&T – growth competition and transition). The subpopulations mutual influence can be either cooperative or suppressive. Please refer to Table 1 for model formalism. **C)** Improvement in goodness of fit (measured by chi-square values) of G&T and GI&T models when cell-state transition rates are allowed to take smaller values with respect to mesenchymal cell growth rates (lower bound on normalised transition rates = 1/(g t ratio); g t ratio: growth-to-transition ratio). **D)** Model selection using Corrected Akaike Information Criteria (AICc) values. As with minimum chi-square values (panel B), GI&T model has lower AICc values than G&T for a given value of g t ratio. At g t ratio of 50, the GI&T AICc is 46.38 while G&T AICc is 49.19. g t ratio = 50 for model fits in panel A. Mesenchymal cell growth rate has been used to non-dimensionalize the model equation and its value is set to (1/50) hrs^-1^ to rescale the time units of model output and compare temporal dynamics with experimental data.

We observed that only a difference in the intrinsic growth rate (model G with E growth rate (r_e_) ≥ M growth rate (r_m_)) could not reproduce the dynamics and eventual 80:20 ratio of EpCAM^high^ and EpCAM^low^ populations when the initial population consists of either M or E cells alone. This inaccuracy led us to further consider a scenario that also accounts for the proliferation rates being dependent on non-cell autonomous effects. Thus, we considered frequency-dependent cooperative or suppressive effect of one subpopulation on growth of the other, besides intrinsic growth differences between E and M subpopulations. Two additional parameters were introduced: σ (the influence of E cells on growth rate of M cells) and μ (the influence of M cells on growth rate of E cells) parameters. However, this Growth Influence (GI) model also failed to adequately explain the observed data (**Figure 1A**). In particular, we saw a sudden change in the fraction of M (EpCAM^low^) cells for GI model around the second week, an observation not seen in experimental data. This sudden change could be attributed to the threshold fraction of E and M populations required for influencing the growth rate of the other subpopulation. Together, these simulations show that a difference in intrinsic growth rate – with or without the influence of one subpopulation on growth rate of the other – alone cannot explain observed experimental data in PMC42-LA breast cancer cells.

We next hypothesized that including cell-state transitions between E and M phenotypes can help explain the experimental data more accurately. Thus, we added cell-state transition terms (E-to-M and M-to-E transitions) to the above models, resulting in Growth and Transition (G&T), and Growth Influence and Transition (GI&T) models with t_em_ and t_me_ as the E-to-M and M-E per-cell transition rates respectively. Both G&T and GI&T gave significantly improved model fits to the experimental data (**Figure 1A**). However, these improvements in model fitting were observed only when the E-M and M-E transition occurred at rates much slower than growth rates of mesenchymal cells (**Figure 1B, S1**). Here, the parameter ‘g-t ratio’ (growth-to-transition rate ratio) determined the lower bound on the search range for E-M and M-E transition rates during the parameter optimization process. The larger the g-t ratio, the smaller the lower bound on transition rates (**Table 3**). Thus, although cell-state transitions are necessary to explain the experimental observation, our findings suggest that they must occur at slower rates relative to cell division to explain the experimentally observed data.

Since, parameter values within their 95% confidence could give equally good fit to the experimental data, we next looked at how model parameters are co-variable with each other. Perhaps intuitively, in order for the G&T model to explain a significant long-run (20%) contribution of M cells in the population, an increase in E growth rates (r_e_) must lead to a concomitant increase in E-M (t_em_) and decrease in M-E (t_me_) transition rates (**Figure S2**). This interdependence between growth and transition rates was subsequently weakened through the inclusion of additional growth-influencing parameters (σ and μ) in the case of the GI&T model (**Figure S3**). In the GI&T model, σ (the influence of E cells on the growth rate of M cells) became more negative with increasing r_e_ values, and thus E cells could increasingly support M cells growth when E cells themselves divided faster. Similarly, the suppression of E cell growth by M cells increased when E cells divided faster (**Figure S3**). Effectively, this addition allows intrinsic growth rates and growth influence parameters to offset one another’s effects, and as a result contributed to unidentifiability of parameters (as discussed in section 2). Growth influence (σ and μ) parameters barely influenced transition rates (t_me_ and t_em_) in most of their 95% confidence range.

We next calculated Corrected Akaike Information Criteria (AICc) values to perform model selection that achieves a trade-off between the goodness of fit and model complexity (number of parameters). We found that in the g-t ratio regime where both the G&T and GI&T models performed their best, GI&T exhibited slightly improved AICc values (**Figure 1C**). These findings suggested that the most probable process that gave rise to the observed experimental phenomenon included :1) intrinsic growth rate differences between E and M cells, 2) cooperative/suppressive growth interactions, and 3) cell-state transitions.

### Cell-state transition, along with frequency-dependent growth influence of subpopulations, explain spontaneous EMP in HCC38 cells

We next focused on another set of EMT population dynamics experiments where co-cultures of different starting ratios of E and M cells were tracked, and the temporal dynamics of phenotypic heterogeneity was recorded over time. In these experiments, the authors co-cultured various percentage of Venus (dye) labelled EpCAM^high^ cells with Venus non-labelled EpCAM^low^ cells, and they tracked increasing fractions of Venus labelled EpCAM^low^ cells in the Venus labelled population over 12 days (Culture 1 to 3 in **Figure 2A**). Similar co-cultures were performed for varied percentage of Venus labelled EpCAM^low^ cells with Venus non-labelled EpCAM^high^ cells to track the increasing fraction of EpCAM^high^ cells (decreasing fraction of EpCAM^low^ cells) in the Venus labelled population over 12 days (Culture 4 and 5 in **Figure 2A**). The experiments demonstrated that the measured fraction of E cells that transitioned to the M state was higher at each measured timepoint when there was a higher initial frequency of M cells in the co-culture population. Similarly, the measured fraction of M cells that transitioned to E cells was higher when there was a higher initial frequency of E cells in the co-culture population. Relatively speaking, the impact of increased percentage of E cells on MET was less than that of increased percentage of M cells on EMT. These experimental data support the idea that interactions between E and M populations can modulate their cell-state transition rates.

**Figure 2.**
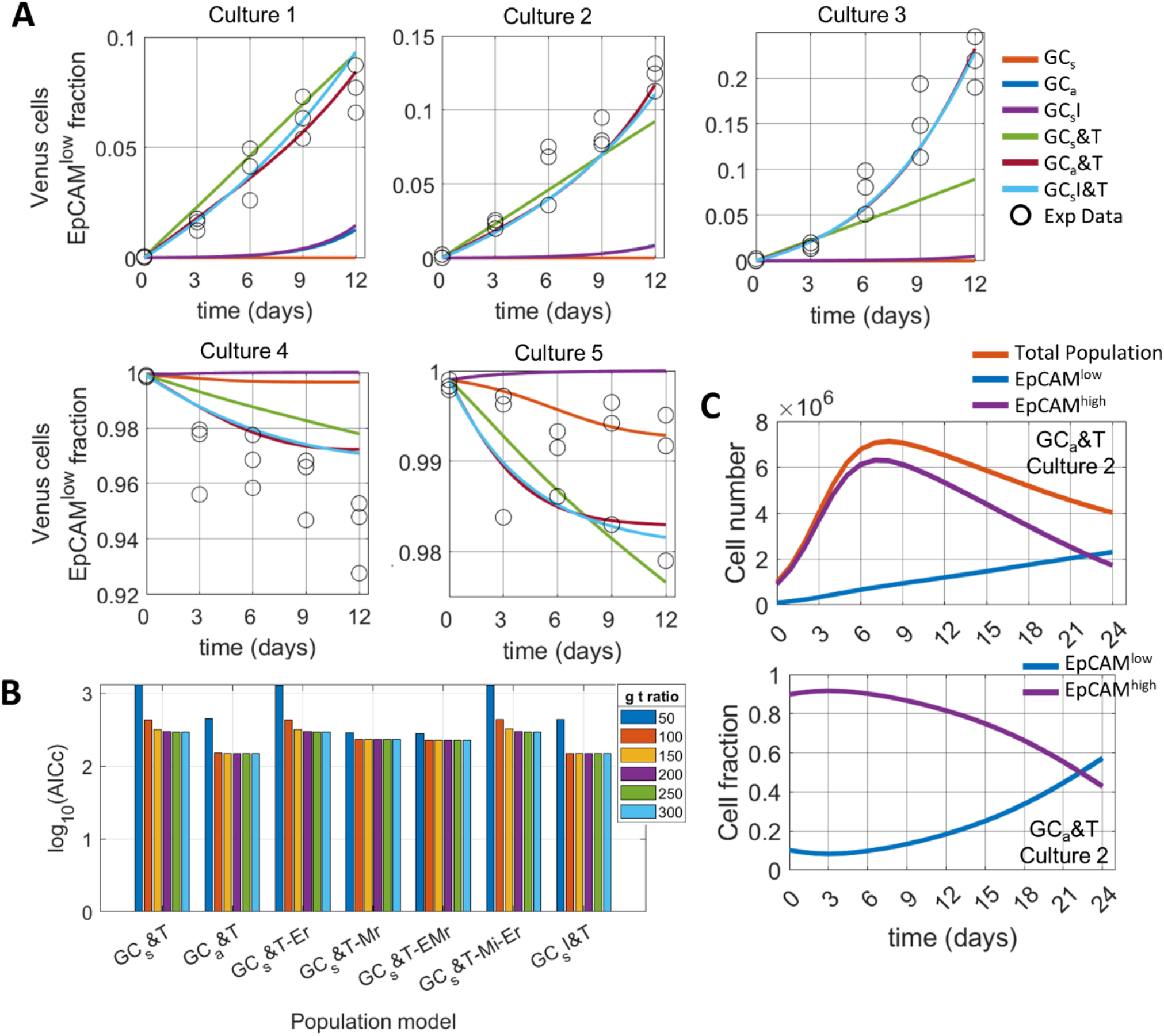
Necessity of both mutual growth competition/influence and cell state transitions among subpopulations to explain spontaneous EMP in HCC38 cells. (Figure 3 in Yamamoto et al. Cancer Science 2017). **A)** Model fits to experimental data of different culture conditions depending upon the initial mixture of Venus label and EpCAM^high/low^ status. The models here capture the following mechanisms/interactions among E and M subpopulations – 1) Symmetric resource competition for growth (GC_s_), 2) Asymmetric resource competition for growth (GC_a_), 3) Symmetric resource competition and growth influence (GC_s_I). We then added cell-state transitions to the above models to obtain GC_s_&T, GC_a_&T and GC_s_I&T models. Please refer to methods and Table 2 for model formalism (The models considered here have logistic growth term unlike those used to fit data in Figure 1, which considered exponential growth). **B)** Comparison of Corrected Akaike Information Criteria (AICc) values across expanded set of models that also include models that consider mutual influence of subpopulations through changes in the transition rates as the population frequency distribution changes with time. Here, ‘Er’ denote that the epithelial cells lower down their E-to-M transition rate with their increasing frequency in the population. Similarly, such retention effect is considered for the M cells as well (GC_s_&T-Mr). Model GC_s_&T-Mi-Er incorporate retention effect of both E and M populations. Term ‘Mi’ in model G&T-Mi-Er indicates that M cells increase the E-to-M transition rates with their increasing frequency in the population. **C)** Temporal dynamics of EpCAM^low^ and EpCAM^high^ cell number and fraction in the population from model GC_a_&T using best fit parameters.

To explain this experimental data, we asked our original question – is it necessary to incorporate cell-state transition in adequately describing the experimental data? The models considered here are analogous to those considered for fitting PMC42-LA cell data, however, we included logistic (instead of exponential) growth term in all the models. We made this change to mimic experimental conditions where the cells were continuously grown without passaging. We considered two types of logistic growth: 1) symmetric competition – where E and M cells consume equal amount of resources, and 2) (two-species ecological) asymmetric competition – where E and M cells consume unequal amount of resources. The experimental study also reported the doubling times of E and M cells to be 35 and 54 hrs, respectively, and therefore the growth rate (r_e_ and r_m_) parameters for all models were set constant as per above values and were not included in the model fitting exercise.

We first considered models that captured only population dynamics due to growth differences, competition and growth influence (cooperative/suppressive) – 1) GC_s_ (logistic growth symmetric competition), 2) GC_a_ (logistic growth asymmetric competition), and 3) GC_s_I (logistic growth symmetric competition with subpopulations influence on growth). However, these models could not explain the experimental data well (**Figure 2A**). Inclusion of cell-state transitions led to significant improvement in the goodness of fit (lower chi-square) values, although again at large g-t ratio (**Figure S4A, B** – model GC_s_&T, GC_a_&T and GC_s_I&T).

To decipher the possible interactions among E-M populations, we expanded our models set to include subpopulation interactions that modulate transition rates apart from growth influence (please refer to **Table 2** and the Methods section for detailed model descriptions). We considered two effects, retention and influence, in the ability of E and M subpopulations to influence cell-state transition rates. By *retention* (denoted by the lower case ‘r’) we mean that a subpopulation reduces its cell-state transition rate to another subpopulation (phenotype) when in majority. On the contrary, the *influence* effect (denoted by the lower case ‘i’) of E/M subpopulations enhances cell-state transition rates of another subpopulation (phenotype) to its own when in majority. The models considered include retention in either E or M cells or both (GC_s_&T-Er, GC_s_&T-Mr and GC_s_&T-EMr), and influence of M cells on E cells (GC_s_&T-Mi-Er). These specific cellular interactions affecting cell-state transitions were considered along the lines of existing reports ^20–22^.

The proposed models were checked for parameter identifiability using *a priori* and *a posteriori* methods ^23–25^. *A priori* structural identifiability of models using a differential algebra approach resulted in all models being structurally unidentifiable in the global parametric ranges. However, *a posteriori* identifiability analysis using the Profile likelihood method resulted in all parameters across models to exhibit well defined upper and/or lower bounds (**Figure S4C, D**). Thus, the data along with the well-defined biologically relevant parameter range (**Table 5**) were together able to resolve parameters in well-defined intervals.

Incorporation of population-specific effects on transition rates did not change the dependence of models incorporating cell-state transitions to require large g-t ratio to adequately fit the data (**Figure S4B**). All the models predict the E-M transition to be much faster than the M-E transition (transition rates t_em_ >> t_me_), which was also reported in the experimental study (**Figure S4A**).

On performing model selection, the use of AICc resulted in model GC_a_&T (which considered asymmetric growth competition with basal cell-state transitions) and GC_s_I&T (which considered symmetric growth competition with influence on growth by subpopulations and basal cell-state transitions) as being the best models with little difference of AICc values among themselves (**Figure 2B**).

Focusing specifically on the parameter co-variability of the GC_a_&T model 95% confidence range analysis using profile likelihood, we observed that growth competition affected the carrying capacity of E and M cells (**Figure S5**). With increasing growth competition to M cells by E cells (α), K_e_ was increased and K_m_ was reduced. The reduced proliferation capacity (K_e_) of E cells was substantiated by more M-E transitions, and less E-M transition with reduced growth competition to E cells by M cells (β) (**Figure S5**). Inversely, increasing growth competition to E cells by M cells (β) led to enhanced proliferation capacity of E cells (K_e_) with concomitant increased E-M transitions, and diminished M-E transitions with reduced growth competition to M cells by E cells (α) (**Figure S5**). Similarly, for GC_s_I&T model, higher growth suppressive effect of M cells on E cells (μ) led to increasing carrying capacity (K), and M-E transition rates; and larger cooperative interaction by E cells (σ) led to reduced E-M transition rates and increased M-E transition rates (**Figure S6**). However, the relations among parameters are based on their co-variability following a best-fit to the data, and so it is therefore intractable to infer causal relations (independent/dependent) among parameters solely from the model fitting exercise.

We next looked at the temporal dynamics of overall EpCAM^low^ and EpCAM^high^ cells, irrespective of their Venus label, for the best fit model GC_a_&T. EpCAM^high^ cell numbers that had grown to large number in the first 10 days of culture started to drop down because of increasing resource competition from the mesenchymal population (**Figure 2C** top panel). These negative growth trends for E cells and positive trends for M cells led to an increasing long-run population fraction of M cells (**Figure 2C** bottom panel). Similarly, even for the GC_s_I&T model, where E cells cooperate/enhance M cells growth and M cells supress E cells growth, M cells dominate the population frequency in long run (**Figure S7**). However, the parental HCC38 cells had an EpCAM^high^:EpCAM^low^ ratio of 90:10. Therefore, we argue that such dominance of EpCAM^high^ cells in the cell line can be reasonably explained by keeping E cells to be in the exponential phase growth, which provides them with an additional advantage over the large growth suppressive influence of M cells and high E-M transition rates.

To test whether the impact of E and M subpopulation frequencies on the cell-state transition rates improved the model fit to experimental data for PMC42-LA cells, we expanded the list of models to include additional terms corresponding to the effects of phenotype on their retention (i.e., E cells prevent their transition to M) and influence on others (i.e., E cells promote MET in M cells) respectively, and fitted them to experimental data (refer **Table 1** and the Methods section for model formalism). Parameter identifiability analysis using analytical differential algebra method results in all models to be globally identifiable conditional on knowing r_m_ and/or r_e_ (SI Text). Consequently, parameter identifiability analysis using an *a posterior* profile likelihood method showed that, given the experimental data and relevant bounds on the parameters (**Table 3**), all model parameters had either well defined upper and/or lower bounds, with the exception of models G&T-Mi and G&T-Mi-Mr, whose transition influence parameters did not have both upper and lower bounds (**Figure S8B, C**). In the model set, model G&T-EMr gave a significantly good fit at smaller g-t ratio values (**Figure S8A**). Since the g-t ratio determines the lower bound on basal transition rates (t_me_ and t_em_), smaller g-t ratio values well-describing the data is understandable given that the added retention effect acts to reduce the effective transition rate (t_me_(1-δ) and t_em_(1-γ)). This allows the G&T-EMr model to attain larger estimates of t_me_ and t_em_ parameters while fitting the data. For larger g-t ratio values, we found that G&T and GI&T had lower chi-square than other models, with GI&T having the lowest AICc values (**Figure S8**). These trends further strengthen that the models incorporating either independent or mutually dependent subpopulation growth rates, together with basal cell-state transition rates, are best to explain the EMP dynamics seen in PMC42-LA cells.

**Table 1.** Model proposed for fitting spontaneous E-M plasticity data seen in PMC42-LA cells

| Model | Description |
| --- | --- |
| G&T | Growth and transition |
| GI&T | Growth Influence and transition |
| G&T-Mr | Growth, transition, and M cells reducing their state transitions |
| G&T-Er | Growth, transition, and E cells reducing their state transitions |
| G&T-EMr | Growth, transition, E and M cells reducing their cell state transitions |
| G&T-Mi | Growth, transition, M cells' influence on E cells for E-M transition |
| G&T-Mi-Mr | Growth, transition, M cells' influence on E cells for E-M transition, and M cells reducing their M-E transition rates |

### Changes in basal cell-state transition rates can explain population dynamics of TGFβ and JAK2/3 inhibitor treatment data in HCC38 cells

To investigate whether cells communicate with each other through paracrine signalling and influence E-M and M-E transition dynamics, Yamamoto *et al*. further used varied EMT inhibitors to inactivate the crosstalk between EpCAM^low^ and EpCAM^high^ cells^9^. They observed that the proportion of cells in the Venus positive EpCAM^low^ state were significantly reduced when TGFβ signalling and JAK2/3 were inhibited relative to control groups.

The abovementioned experimental findings suggested that cell-cell communication impacted cell-state transition rates. However, even though our model set included models with the impact of cell-cell communication on the transition rates, our previous analysis suggested that the GC_a_&T and GC_s_I&T models gave the best fit for the spontaneous E-M plasticity data. Thus, we investigated whether changes in cell-cell communication via modulation of growth or transition rates improves the fit to the experimental data related to TGFβ and JAK2/3 inhibitor treatment. An alternate scenario can be that treatment with these inhibitors changes the basal cell-state transition rates. To assess these possibilities, we considered following three scenarios: 1) Optimizing for basal transition rates (t_me_ and t_em_) with inhibitor treatment data while taking best estimates of other parameters (growth/transition influence and carrying capacity) from analysis on spontaneous data, 2) Optimizing for growth/transition influence parameters with inhibitor treatment data while taking best estimates of other parameters (transition rates and carrying capacity) from analysis on spontaneous data, and 3) Optimizing for both basal transition rates and growth/transition influence parameters with inhibitor treatment data while obtaining best estimates of other parameters (carrying capacity) from analysis on spontaneous data. Although the lowest normalised chi-square value (normalised by number of estimated parameters) across models for each inhibitor treatment resulted from scenario 3, the lowest AICc values were obtained for scenario 1 (**Figure 3B, Figure S9**). We therefore conclude that inhibitor treatment modulated the basal transition rates more significantly than the cell-cell communication (via growth/transition influence parameters), leading to reduced EMT (**Figure 3A**). Also, models GC_a_&T and GC_s_I&T had either comparable or significantly lower AICc values with respect to other models for the scenario 1 to explain the inhibitor treatment data. Collectively, these findings further assured us that the asymmetric growth competition and growth influence model with basal transitions (GC_a_&T and GC_s_I&T models) are the most probable mechanisms that underlie spontaneous and EMT inhibitor-induced dynamics seen in **Figure 2A** and **Figure 3A**.

**Figure 3.**
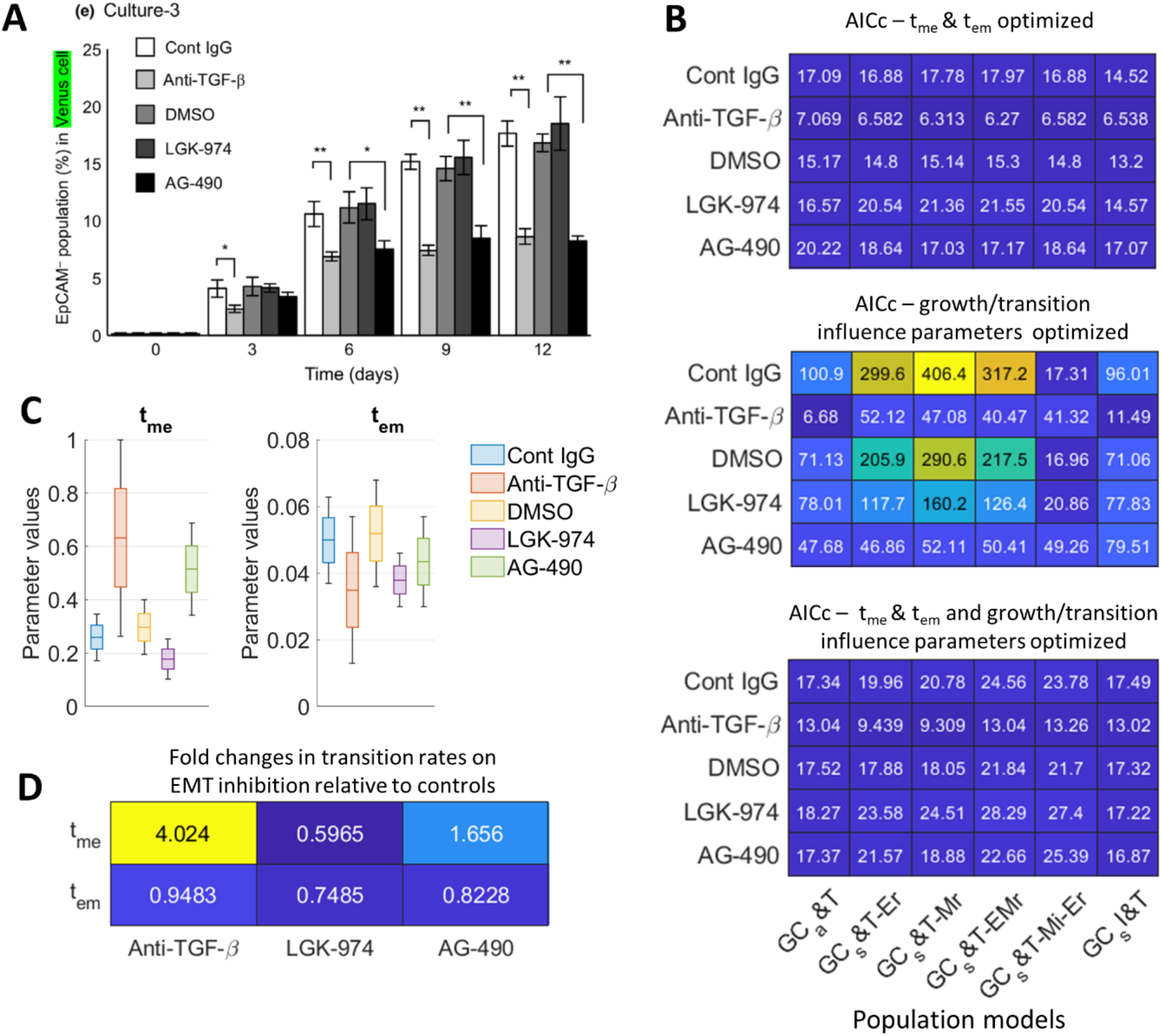
Changes in basal transition rates better explain the EMT inhibition data than changes in cell-cell communication effects. **A)** Effect of EMT inhibitors on E-M cell-state transitions (adopted from Yamamoto et al, Cancer Science 2017). Here, Cont IgG is the control case for Anti-TGFβ, and DMSO is the control case for both KGK-974 and AG-490. **B)** AICc values of each model upon fitting to the differentinhibitor treatment data. Top (resp. middle) panels correspond to scenarios where only basal transition rates (resp. influence parameters) were optimized and the remaining parameters were estimated from the previous analysis of Figure 2 and held constant. Bottom panel corresponds to AICc values when both basal transition rates and influence parameters were optimized. **C)** Distribution of the basal transition rates in GC_a_&T model across different inhibitor treatment data. **D)** Fold change in the best fit t_me_ and t_em_ parameters of GC_a_&T model for the inhibitor treatment data relative to their controls.

We next looked at the changes in the basal transition rates t_me_ and t_em_ resulting from the inhibitor treatments (**Figure 3C-E**). The t_me_ rates were significantly larger for post treatment with Anti-TGFβ and AG-490 molecules than for their respective controls (Control IgG and DMSO). On the other hand, t_em_ was reduced, but not significantly for both TGFβ and AG-490 inhibition case (**Figure 3C**). When comparing the best-fit t_me_ and t_em_ parameters between controls and inhibitor treatment data fitting, we saw 4- and 1.5-fold increases in t_me_ values on treatment with Anti-TGFβ and AG-490, respectively, with only slight decrease in the t_em_ values (**Figure 3D**). Therefore, we conclude that increased M-E transition rates contribute to reduced EMT following inhibitor treatment.

### Analysing model selection uncertainty via uniqueness of temporal dynamics and sensitivities to experimental data

The best fit model need not necessarily be closest to the underlying cellular mechanism generating the observed dynamics, given the coupled impact of noise in cellular process with additional noise in experimental measurement. Thus, to quantify how much uncertainty lies in model selection for noisy data as well as what specific experimental conditions a given model is most sensitive to, we performed the following two analyses: 1) Cross-model fitting, and 2) Cross-one validation (leave-one-out) – for models proposed to fit HCC38 cells data.

In cross-model fitting, we checked whether the temporal dynamics resulting from a model, given a parameter set, could be well-explained by any other model in the list (**Table 2**). For this, synthetic data sets (having identical structure to the original experimental setup) were generated using randomly sampled parameters for all models in Table 2. For each synthetic data set generated from a known underlying mechanism (y axis in **Figure 4A**), every model was used to fit the synthetic data (x axis in **Figure 4A**), and the procedure was repeated for a total of 1000 synthetic data sets overall, corresponding to 1000 independent, randomly sampled parameter sets for each model. After fitting synthetic data from each model to all the models, the AICc criteria was used to identify a model that strikes a trade-off between the goodness of fit and model complexity (number of parameters). Each row of the heatmap (shown in **Figure 4A**) denotes the percentage distribution of models on the x-axis selected by AICc criteria as best model (which has the highest probability to have generated the data) while fitting to the 1000 synthetic data from a model on y axis. We found that all models performed poorly for being selected as the best model when they fitted other models generated synthetic data with a small amount of noise. However, the exception to this trend is the GC_s_&T-EMr model, which could well explain the data generated for cell-state transition influence models (Er, Mr, and Mi-Er) well and that generated for the growth modulation scenario (GC_s_I&T) (refer to columns of **Figure 4A,i**). However, increasing the level of noise in the synthetic data, in general, led to: 1) higher rates of model misidentification (false positives), which was especially true of GC_s_&T and GC_a_&T models as they were selected as the best-fit model for data which they did not generate (columns corresponding to GC_s_&T vs. GC_a_&T in **Figure 4Ai, ii**); and 2) lower rates of correct identification (true positives), wherein the selected best-fit models had actually generated the synthetic data (diagonals of two heatmaps, **Figure 4A**). Thus, increasing noise in the experimental data blurred our framework’s ability to identify the underlying mechanism generating the data.

**Table 2.** Model proposed for fitting spontaneous E-M plasticity data seen in HCC38 cells

| Model | Description |
| --- | --- |
| GC<sub>s</sub>&T | Logistic growth (symmetric competition), and transition |
| GC<sub>a</sub>&T | Logistic growth (asymmetric competition), and transition |
| GC<sub>s</sub>&T-Er | Logistic growth (symmetric competition), transition, and E cells reducing their state transitions |
| GC<sub>s</sub>&T-Mr | Logistic growth (symmetric competition), transition, and M cells reducing their state transitions |
| GC<sub>s</sub>&T-EMr | Logistic growth (symmetric competition), transition, and both E and M cells reducing their state transitions |
| GC<sub>s</sub>&T-Mi-Er | Logistic growth (symmetric competition), and M cells' influence on E cells for E-M transition, and M cells reducing their M-E transition rates |
| GC<sub>s</sub>I | Logistic growth (symmetric competition) and growth influence of E and M subpopulations on each other |
| GC<sub>s</sub>I&T | Logistic growth (symmetric competition), growth influence of subpopulations, and cell-state transitions |

**Table 3.** Parameter search ranges while optimizing models proposed for fitting PMC42-LA E-M heterogeneity emergence data

| Parameter | Description | Search range |
| --- | --- | --- |
| $r_m$ | M cell growth rate (1/hr) | $r_m$ |
| $r_e$ | E cell growth rate (1/hr) | $[r_m \ 3r_m]$ |
| $t_{me}$ | M-E transition rate (1/hr) | $[\frac{r_m}{g \ t \ ratio} \ r_m]$ |
| $t_{em}$ | E-M transition rate (1/hr) | $[\frac{r_m}{g \ t \ ratio} \ r_m]$ |
| $\sigma$ | E cells influence on M cells growth rate | $[-1 \ 1]$ |
| $\mu$ | M cells influence on E cells growth rate | $[-1 \ 1]$ |
| $\gamma$ | strength of E cells to reduce E-M transition | $[0 \ 1]$ |
| $\delta$ | strength of M cells to reduce M-E transition | $[0 \ 1]$ |
| $\theta$ | strength of M cells to enhance E-M transition | $[1 \ 4]$ |

**Table 4.**
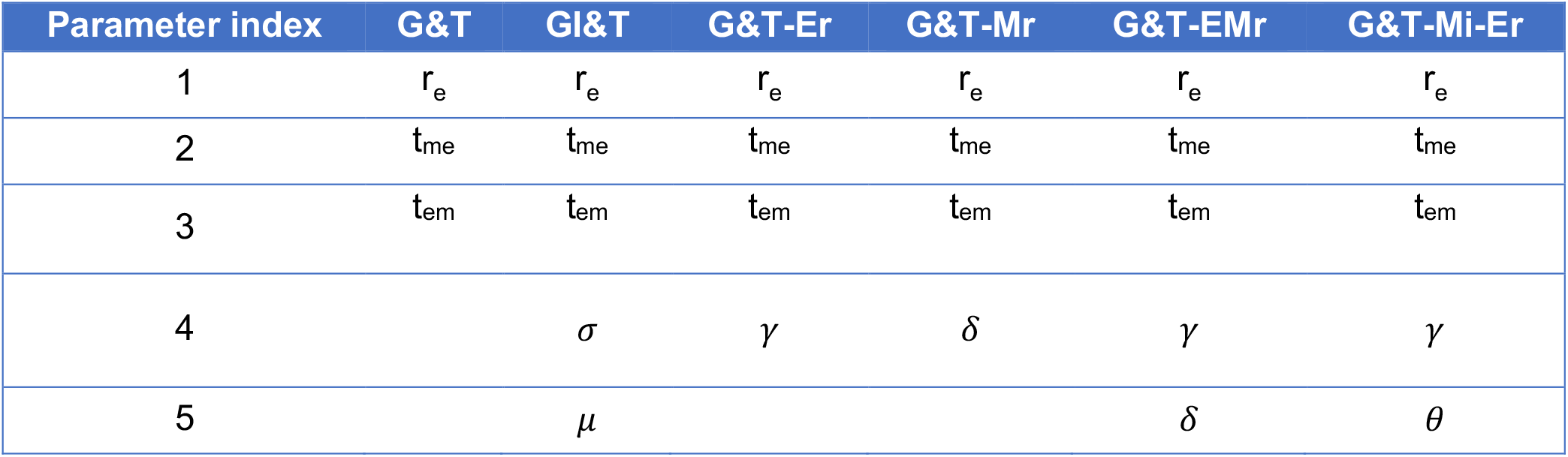
Model wise parameter list (corresponding to models proposed for fitting PMC42-LA E-M heterogeneity emergence data).

**Table 5.** Parameter search ranges while optimizing models proposed for fitting PMC42-LA E-M heterogeneity emergence data.

| Parameter | Description | Search range |
| --- | --- | --- |
| $r_m$ | M cell growth rate (1/hr) | $r_m$ |
| $r_e$ | E cell growth rate (1/hr) | $r_e$ |
| $t_{me}$ | M-E transition rate (1/hr) | $[\frac{r_m}{g\ t\ ratio} \ r_m]$ |
| $t_{em}$ | E-M transition rate (1/hr) | $[\frac{r_m}{g\ t\ ratio} \ r_m]$ |
| $\alpha$ | E cells influence on M cells growth rate | [0 10] |
| $\beta$ | M cells influence on E cells growth rate | [0 10] |
| $\gamma$ | strength of E cells to reduce E-M transition | [0 1] |
| $\delta$ | strength of M cells to reduce M-E transition | [0 1] |
| $\theta$ | strength of M cells to enhance E-M transition | [1 4] |
| $\sigma$ | E cells influence on M cells growth rate | [-1 1] |
| $\mu$ | M cells influence on E cells growth rate | [-1 1] |
| K | Common carrying capacity (cells) | $[10^6 \ 20 \times 10^6]$ |
| $K_e$ | E cells carrying capacity (cells) | $[10^5 \ 20 \times 10^6]$ |
| $K_m$ | M cells carrying capacity (cells) | $[10^5 \ 20 \times 10^6]$ |

**Table 6.** Model wise parameter list corresponding to models proposed for fitting PMC42-LA E-M heterogeneity emergence data.

| Parameter index | G&T | GC<sub>a</sub>&T | G&T-Er | G&T-Mr | G&T-EMr | G&T-Mi-Er | GC<sub>s</sub>I | GC<sub>s</sub>I&T |
| --- | --- | --- | --- | --- | --- | --- | --- | --- |
| 1 | $t_{me}$ | $t_{me}$ | $t_{me}$ | $t_{me}$ | $t_{me}$ | $t_{me}$ | $t_{me}$ | $t_{me}$ |
| 2 | $t_{em}$ | $t_{em}$ | $t_{em}$ | $t_{em}$ | $t_{em}$ | $t_{em}$ | $t_{em}$ | $t_{em}$ |
| 3 | K | $\alpha$ | K | K | K | K | $\sigma$ | K |
| 4 | | $\beta$ | $\gamma$ | $\delta$ | $\delta$ | $\gamma$ | $\mu$ | $\sigma$ |
| 5 | | $K_e$ | | | $\gamma$ | $\theta$ | K | $\mu$ |
| 6 | | $K_m$ | | | | | | |

**Figure 4.**
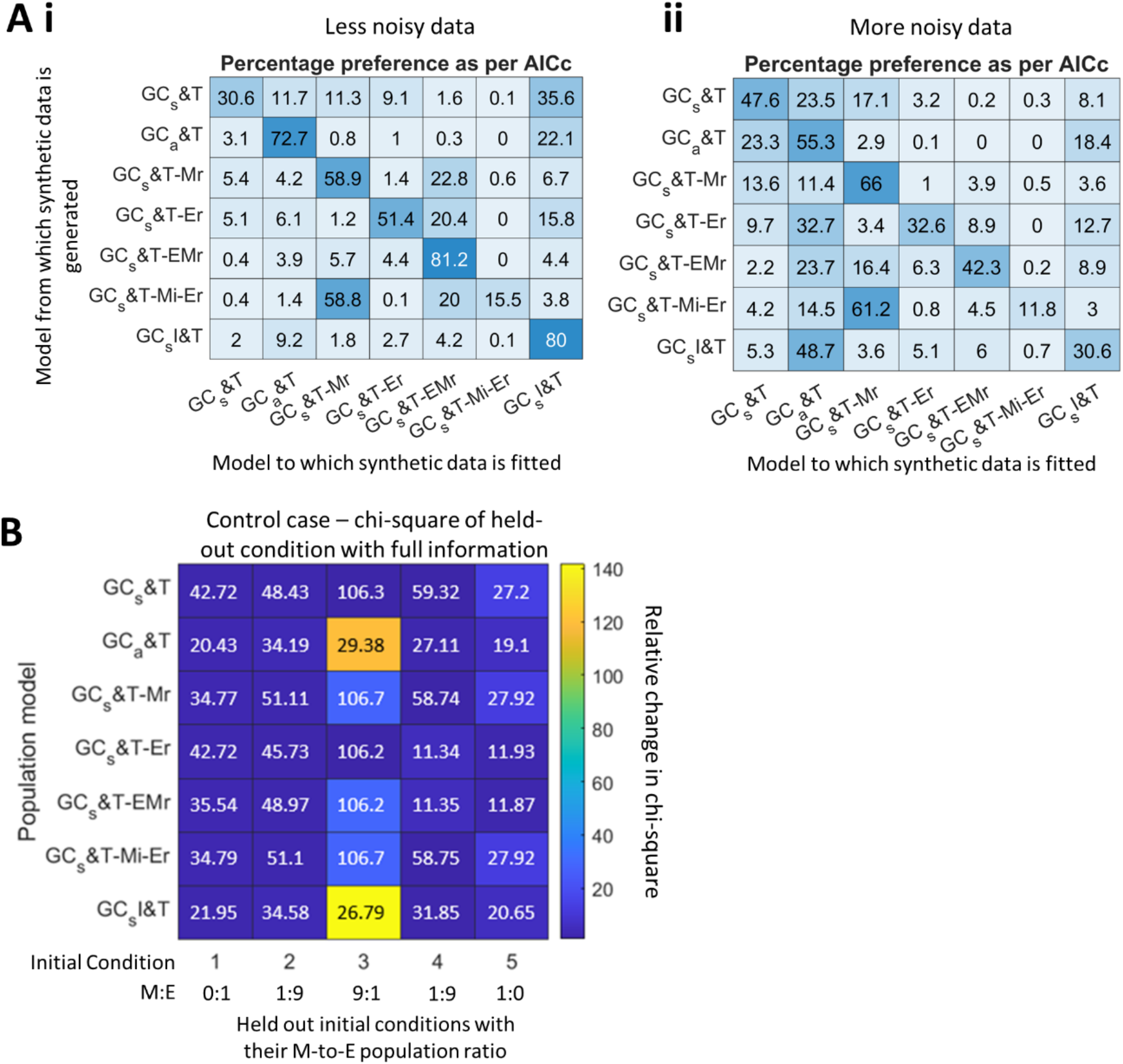
Analyzing model selection uncertainties via uniqueness of temporal dynamics and sensitivities to experimental data. **A)** Synthetic data was generated using randomly sampled parameters from each model (y axis) and then all the models were made to fit the synthetic data (x axis). The procedure was repeated for 1000 synthetic data sets overall for each model. The values shown in each box of the heatmap denotes the percentage of times a model on the x-axis was selected by AICc criteria as the best model while fitting to the 1000 synthetic data from the models on y-axis. **B)** Leave-1-out analysis. Each culture condition was withheld from fitting, one at a time, and then goodness of fit was estimated using the model against the holdout culture condition. The resulting goodness of fit was then normalized by the control case where goodness of fit (chi-square values) to the same held-out culture condition was calculated when all culture conditions (including the one held out) were used to estimate the parameters. The above calculated relative change in goodness is shown in the colour map, and the numerical values in each block denotes the control case goodness of fit (chi-square) values. For leave-one-out analysis, r_m_ = (1/54) hrs^-1^, r_e_ = (1/35) hrs^-1^ and g t ratio = 250; analysis in panel A is agnostic to absolute parameter values if same values/ranges are used for all model’s data generation and fitting.

Next, in cross-1-validation analysis, the goal was to check sensitive a model is to initial phenotypic distribution of E and M cells in terms of estimating parameters and recapitulate E-M population dynamics. For performing this analysis, each culture condition (in panel A) of the experimental data was withheld, one at a time, while fitting the model and estimating the model parameters. The goodness of fit (chi-square values) was then calculated for the holdout culture condition. This fit value was normalized by the control case where goodness of fit (chi-square values) to the same holdout culture condition was calculated when all culture conditions (including the holdout) were used to estimate the parameters. In **Figure 4B**, the colour bar denotes the normalised chi-square values for each holdout condition, and the numerical values denote chi-square values of the respective control case. Here, we see that although the relative change in goodness of fit is most significant for GC_s_I&T and GC_a_&T models with culture condition 3 (initial M-to-E population ratio, M:E = 9:1), only the above models gave a good fit to culture 3 in their respective control cases. Therefore, even if the models GC_s_I&T and GC_a_&T were significantly more sensitive to culture condition 3, the absolute change in chi-square values across models can be of similar order. All models were relatively less sensitive to other culture conditions (relative change < 20) while also giving good fit to them in their respective controls. Thus, from a modelling standpoint, it is more informative to have experimental data on culture conditions similar to culture 3 (initial 10% E and 90% M proportion) to delineate GC_a_&T or GC_s_I&T from other models and further strengthen the conclusion that one of these models capture the true cellular mechanism giving rise to E-M state plasticity.

Since, in both the analyses above, either generated synthetic data with structure similar to the experimental data or we used partial experimental data, performing these analyses for very limited PMC42-LA breast cancer data could themselves introduce biases. Therefore, we haven’t reported the results for above two analyses for the PMC42-LA models.

### Adding new experimental data points using the best fit model to improve confidence bounds on parameters

The best fit model for HCC38 cell line experimental data (GC_a_&T) had two practically unidentifiable parameters – β (growth competition to E cells by M cells) and K_e_ (carrying capacity of E cells) – with an associated poorly defined upper bound and 95% confidence range (**Figure 5Ai**). For every value of β and K_e_ below the profile likelihood plot threshold (dashed black line), other model parameters were optimized such that the goodness of fit did not change. This insensitivity of the chi-square values for β and K_e_ parameter variation propagate to the invariability of temporal trajectories of Venus cells EpCAM^low^ fraction for the optimized parameter sets. However, such invariability of temporal trajectories to parameter variation is not case for the other practically identifiable model parameters (**Figure S10**).

**Figure 5.**
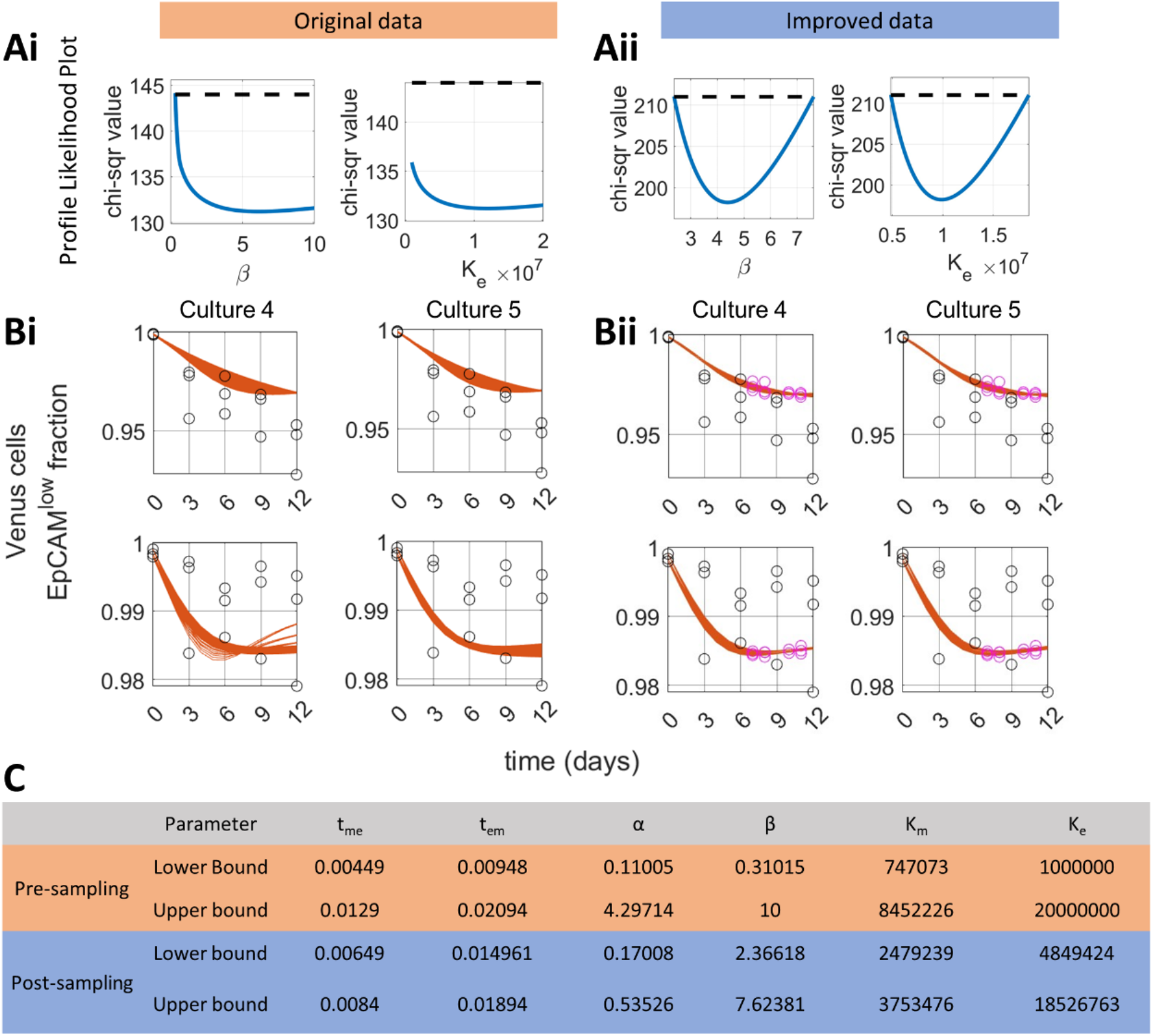
Adding new, model estimated, experimental data points to improve 95% bounds on parameters. **A)**Profile likelihood plots using original and improved data for parameters β and K_e_. **B)** Sensitivity analysis for the dynamics of the Venus EpCAM^low^ cell fraction under variation in β and K_e_ values (constrained by threshold (dashed black line) in panel Ai and Aii, respectively; newly added data points shown in magenta in panel Bii). **C)** Improved 95% confidence range of all model parameters post addition of new data points to experimental data.

To improve on the parameter bounds of β and K_e_, we adopted a method discussed by *Rau et al. 2009*, which identifies a time point of maximum variability in the output (experimental) variable with respect to parameter of interest and adds new (model-estimated) data corresponding to that time point, and then re-runs the parameter identification analysis on the improved data to check for confidence bound improvement^24^. The overall idea is to improve the characteristics of the existing data to better estimate the model parameters by the addition of new data at the informed timepoint.

We implemented this algorithm iteratively, i.e., adding a newer data point if previous addition did not improve the bounds on β and K_e_ parameters until a limit of four additional data points was reached (**Figure 5B,ii**). Given that the output variable (Venus EpCAM^low^ cell fraction) only slightly varied with considerable change in β and K_e_ parameters, the addition of new data points must be accurate (variation among experimental replicates should be small enough) so that even the slight variation in the output with respect to above parameters is captured. Therefore, we saw that the added new data points had very little variation among the replicates (**Figure 5B ii** – circles in magenta at each new time points). The iterative addition of four new data points led to a well-defined upper bound for both β and K_e_ parameters while also further constraining the 95% confidence range of other model parameters (**Figure 5A ii, 5C**).

## Discussion

Recent experimental and computational analyses have investigated the dynamics of spontaneous and drug-induced cell-state transitions. By fitting a Markov chain model to longitudinal data on the proportion of breast cancer stem and non-stem cell subpopulations, the non-stem-to-stem transition was found to be necessary to explain the experimental data ^26^. Similar conclusions were drawn even after accounting for growth rate differences among subpopulations ^15^. Cell-state transitions have also been reported among drug-sensitive and drug-resistant cells, wherein rare drug-resistant cell types either exist prior to treatment or cells acquire resistance over the duration of treatment on short-time scales ^2,27–31^. It becomes particularly difficult to deconvolute the relative contribution of cell division rates and cell-state transitions in experimental settings that occur over timescales spanning multiple cell divisions. Thus, identifying the mechanisms underlying experimental time-course data on the relative frequencies of different interconvertible phenotypes in a cell population remains a formidable task. Nonetheless, mathematical models incorporating cell-state transitions have been able to fit experimental data adequately, thus reinforcing our observations ^29,32,33^.

Our computational model predicts that for HCC38 breast cancer cells, another mode of interaction among the two subpopulations – asymmetric resource competition – is also important. While this prediction remains to be experimentally tested in these cells, resource competition between drug-sensitive and drug-resistant cells is the fundamental idea underlying adaptive therapy that has already shown promise in delaying the occurrence of resistant in several clinical trials ^34^. Furthermore, different tumour clones or phenotypes need not always compete with one another for resources; they may also reciprocally facilitate growth such that their mutual gain leads to highly aggressive and metastatic disease^35,36,38^. Thus, identifying the nature of interaction between different subpopulations within a tumour is crucial to design an optimal treatment strategy. However, little is known about how tumor epithelial and mesenchymal cells interact in distinct (resource-rich vs. resource-limited) micro-environments. Answering this question is especially pertinent given the association between E-M phenotypic states and other axes of cell-state plasticity, such as drug-resistance, stemness and metabolic reprogramming ^37,39,40^. In our analysis, the population dynamics of HCC38 cells is best explained by the GC_a_&T and GC_s_I&T models. Our results for the GC_a_&T model demonstrate that the M cells exhibit high resource consumption with a carrying capacity an order of magnitude smaller than that of E cells. Further, M cells offer a higher competitive pressure for growth of E cells, as compared to *vice versa* (growth competition parameters β > α). Thus, we find that, in the resource limited setting (logistic growth), the asymptotic distribution of the population tends to be M cell-predominant (**Figure 2C**). Therefore, the predominance of E cells in the HCC38 cell line could only be explained if E cells are always maintained in the exponential phase of growth.

Our model also predicts that EMT inhibitors have a larger effect on changes in basal transition rates than they do on cell-cell communication. This could have occurred through two possible ways: 1) modulating only cell-cell communication effects were insufficient to give the overall fold-change required in the effective transition rates (t_me_ and t_em_) to explain the data well of inhibitor treatment. In our models, the cell-cell communication factor changing the effective M-E transition were M cell’s phenotype retention (δ) only, while factors changing E-M transition were E cell’s retention (γ), and M cells influence on E cells (θ). 2) The inhibition of EMT, on blocking of TGFβ with anti-TGFβ antibodies, could help E cells to proliferate faster as EMT induction can suppress growth^19^. Therefore, a selective (growth) advantage may have shifted the population distribution favouring E cells in the experiment, but since the growth rate of E cells was kept constant during our data fitting exercise, the required parametric changes to explain the data well might have reflected into basal transition rates. Additional co-culture experimental data with and without EMT inhibitor data – in the form of greater numbers of initial culture conditions, temporal data points, or direct experimental estimation of cell growth rates – is necessary to confirm above two hypotheses.

The larger ratio of total numbers of experimental data points to the model parameters usually results in well-defined confidence bounds on parameter values. However, if the data does not contain information of certain cellular processes, then even having sufficient time points at which the data is sampled cannot help in bounding parameter values. For example, in the G&T model proposed for fitting E-M transitions in PMC42-LA cells, *a priori* model identifiability analysis predicts that growth rates of E and M cells are not simultaneously identifiable, and this is specific to model cell fraction dynamics (Supplementary file – Structural identifiability analysis). Therefore, we need to set either of the growth rates to a constant value to obtain well-defined estimated bounds of the other growth parameter. Thus, the measured variables in the experimental data determine the identifiability of proposed model parameters. It is for this reason that the only measurement of Venus cell EpCAM^low^ fraction over time in HCC38 cells did not yield well-defined bounds on growth competition parameter (β) and the carrying capacity (K_e_) of GC_a_&T model. Consequently, our efforts to improve bounds on β and K_e_ in **Figure 5** required very precise measurement of Venus cell EpCAM^low^ fraction at newly added time points. These precise measurements helped capture variability in the output resulting from the above two parameters (β and K_e_) variation. Extensive computational platforms are being developed for informing future experimental design to ensure that the best set of system variables are captured at the precise timepoints and model parameter estimates from the data have well-defined confidence bounds ^41–43^.

Our study has many limitations. First, limited experimental data presents a difficulty to construct a complex model giving insights into both growth competition and modulation of cell-state transitions contributions to E-M population dynamics. Second, the population-level mathematical models explored here do not exhaustively capture all conceivable mechanisms by which cells can interact with each other. For example, the assumption that E and M fractions modulate growth and/or transition rates in frequency dependent manner requires well-mixing amongst the cell populations and ignores local interactions between neighbouring cells that can impact spatial heterogeneity in EMT ^20^. The well-mixed case assumption also neglects the concentration gradient of growth factors /cytokines resulting from tissue diffusion. Third, the results on E-M population dynamics presented here may deviate from *in vivo* environments where E and M cancer cells compete with other cell-types present in the microenvironment that also modulate E-M state transition rates. Nonetheless, with limited experimental data and different assays, we were able to narrow down to mechanisms that could explain E-M population dynamics observed in PMC42-LA and HCC38 cells, and establish the necessity of cell-state transition in determining these dynamics. Future longitudinal experimental data using more than one surface or molecular marker to classify phenotypic heterogeneity (for instance, E-cadherin and Vimentin) will facilitate characterizing and isolating the more plastic hybrid E/M phenotypes and their contribution to population dynamics^5,13,44–48^.

## Materials and Methods

### Models

**Table 1** represents the models that are proposed to fit spontaneous EMP seen in PMC42-LA breast cancer cells^12^. All models consider exponential cell growth, with growth rate of the M subpopulation (r_m_) set to a constant value and all rate parameters (E-growth, E-M and M-E transition rates) being scaled by M cells growth rates at the time of parameter optimization. Further, we constrained the growth rate of the E subpopulation (r_e_) to be greater than that of M subpopulation (r_m_) as reported experimentally ^17,18^, giving a normalised r_e_ ≥ 1. The temporal dynamics is realised in terms changes in E and M cell fractions in the population, rather than changes in E and M cell numbers. Following are the mathematical description of each model:

### Growth (G)

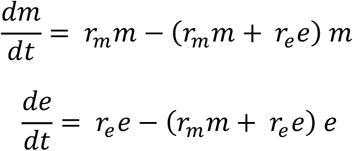

Where, m and e are M and E cell fraction in the population, and r_m_ and r_e_ are the M and E cells growth rates, respectively. The product of (*r*_*m*_*m*+*r*_*e*_*e*) term with *m* in first equation and with *e* in the second equation above comes because of the conversion of cell number dynamics to cell-fraction dynamics.

### Growth Influence (GI)

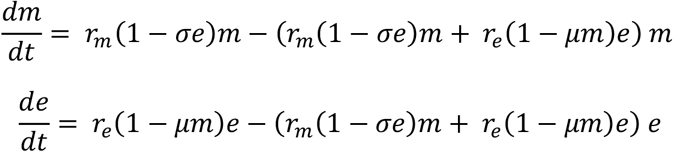

Where, σ and μ are the influence of E cells on M cells’ growth and M cells influence on E cell’s growth. The range of σ and μ during parameter optimization is set between [-1, 1].

### Growth and transition (G&T)

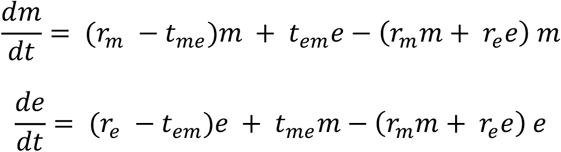

Where, t_me_ and t_em_ are the per-capita M-E and E-M cell-transition rates.

### Growth Influence and Transition (GI&T)

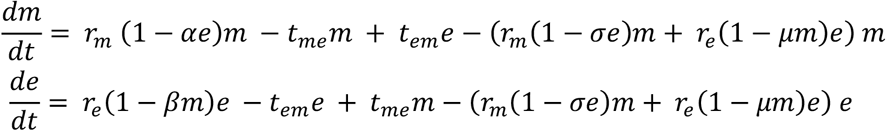

### Growth and Transition – Epithelial retention (G&T-Er)

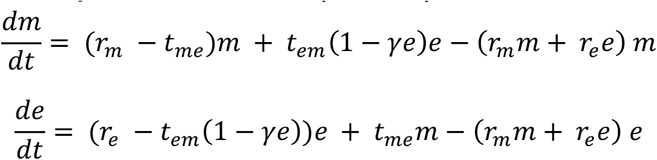

Where, γ sets the strength of E cells to reduce their E-M transition rates. The range of γ is set between [0,1] during optimization.

### Growth and transition – Mesenchymal retention (G&T-Mr)

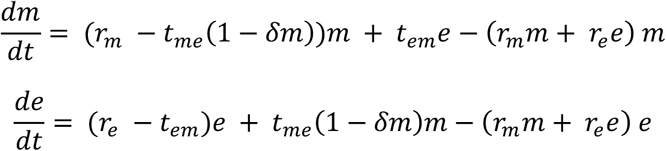

Where, parameter δ sets strength of M cells to reduce their M-E transition rates. The range of *θ* is set between [0,1] during optimization.

### Growth and Transition – Epithelial retention (G&T-MEr)

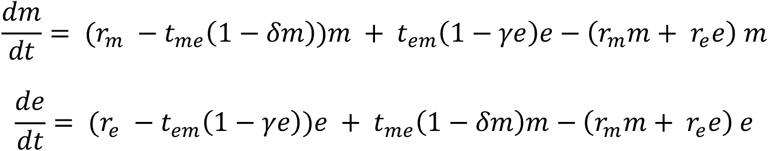

### Growth and Transition – Mesenchymal influence and Epithelial retention (G&T-Mi-Mr)

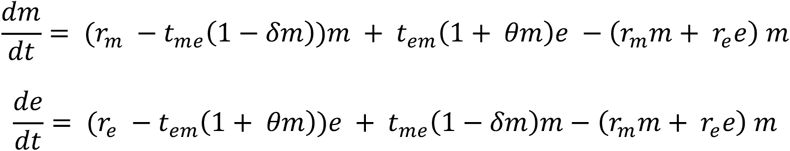

Where, *θ* sets the strength of M cells to enhance the E-M transition rates. The range of *θ* is set between [1,4] during optimization.

**Table 2** represents the models proposed to fit spontaneous EMP seen in HCC38 breast cancer cells^9^. All models consider logistic growth of cells, with all rate parameters (E-growth, E-M and M-E transition rates) being scaled by M cells growth rates at the time of parameter optimization. Further, the temporal dynamics is realised in terms of changes cell numbers of four subpopulations – 1) Venus non-labelled EpCAM^low^ cells, 2) Venus non-labelled EpCAM^high^ cells, 3) Venus labelled EpCAM^low^ cells, and 4) Venus labelled EpCAM^high^ cells. The characteristics (growth, resource competition, transition, and cell-cell influence) of Venus labelled and non-labelled population for a given EpCAM status are kept same. Following are the mathematical description of each model:

### Growth (GC_s_)

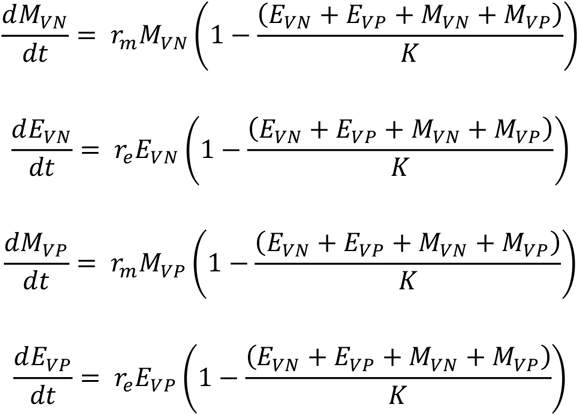

Where, E_VP/VN_ and M_VP/VN_ are EpCAM^high^ and EpCAM^low^ cell numbers with subscript VN and VP denoting their Venus positive and Venus negative status. r_e_ and r_m_ are growth rates of E (EpCAM^high^) cells and M (EpCAM^low^) cells, respectively. K is the carrying capacity (maximum population size). Total E cells, E = E_VN_ + E_VP_, and total M cells, M = M_VN_ + M_VP_.

### Growth and Transition (GC_s_&T)

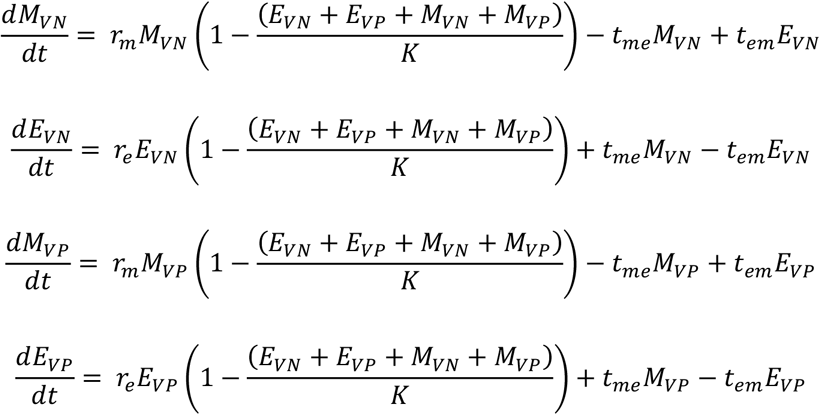

Where, t_me_ and t_em_ are the per-capita M-E and E-M cell-transition rates.

### Growth (GC_a_)

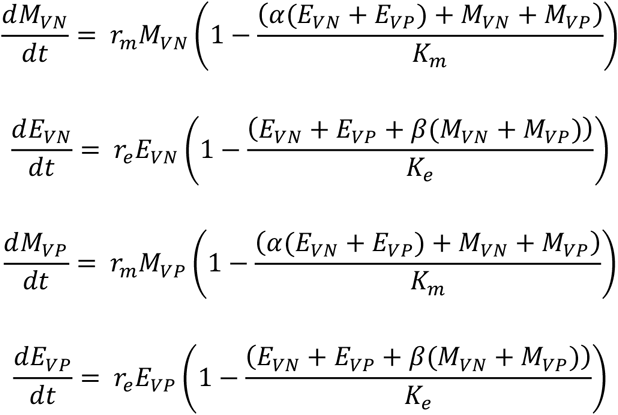

Where, α and β parameters capturing E cell’s influence on M cells growth and M cell’s influence on E cell’s growth. The range of both α and β parameters is in [0 10] during the parameter optimization. K_e_ and K_m_ are the respective carrying capacity of E and M cells.

### Growth and Transition (GC_a_&T)

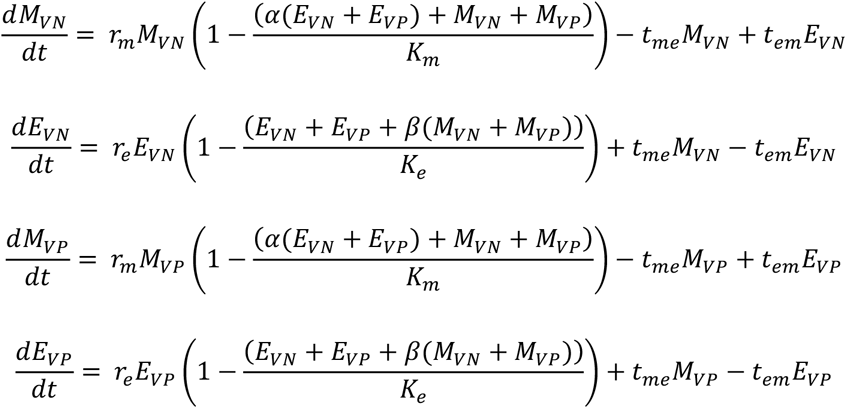

### Growth and Transition (GC_s_&T-Er)

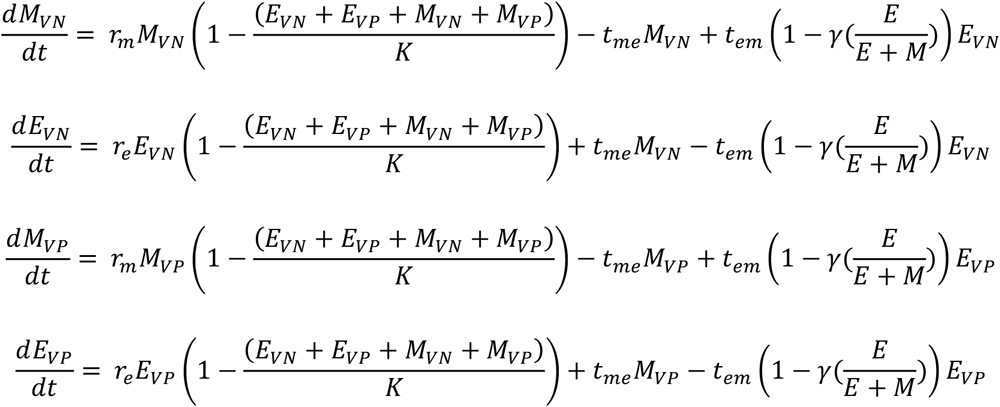

Where, γ sets the strength of E cells (E = E_VN_ + E_VP_) to reduce their E-M transition rates. The range of γ is set between [0,1] during optimization.

### Growth and Transition (GC_s_&T-Mr)

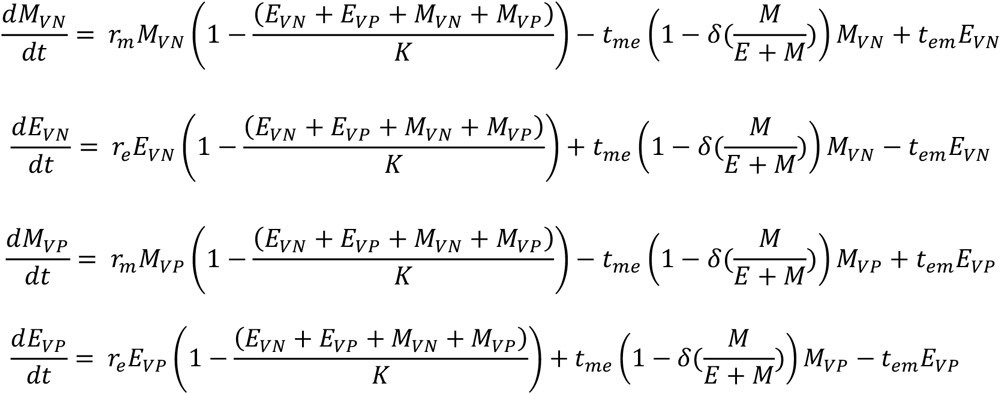

Where, δ sets the strength of M cells (M = M_VN_ + M_VP_) to reduce their M-E transition rates. The range of δ is set between [0,1] during optimization.

### Growth and Transition (GC_s_&T-EMr)

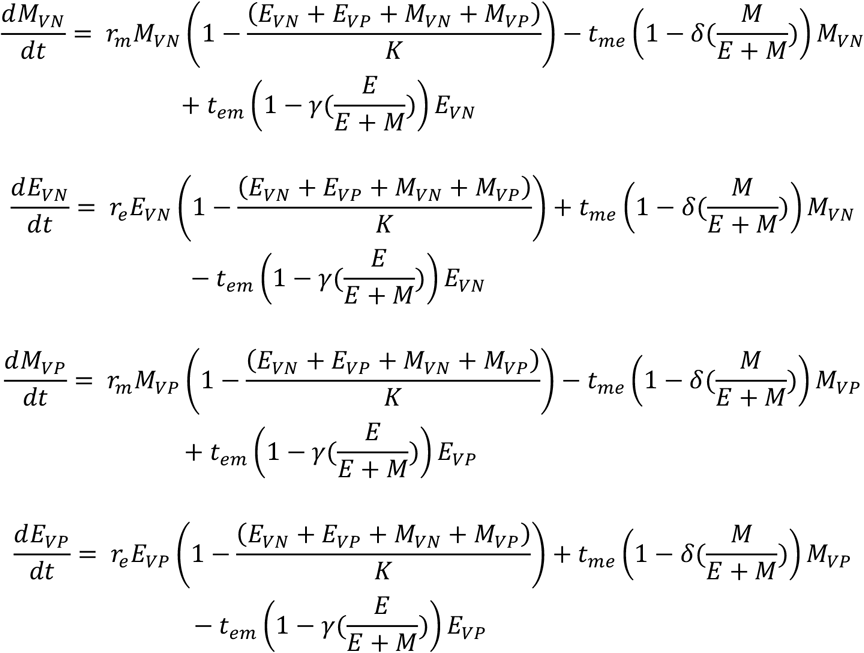

### Growth and Transition (GC_s_&T-Mi-Er)

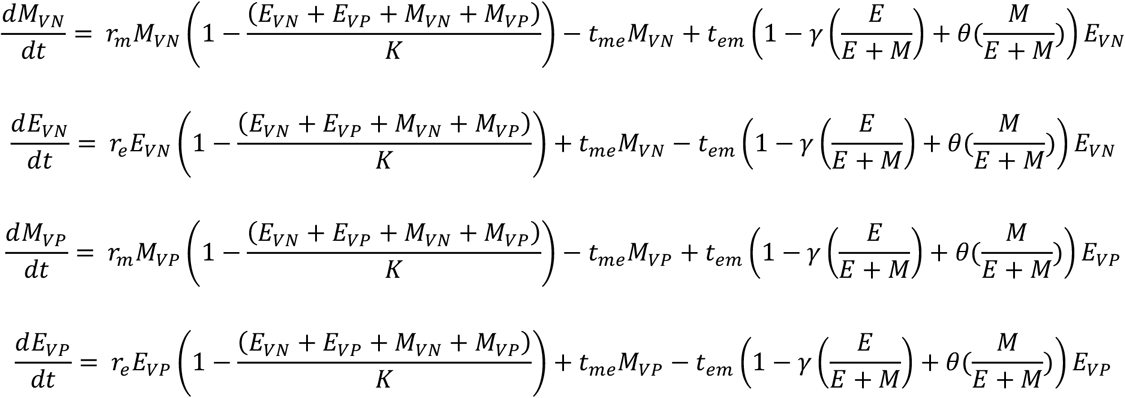

Where, θ sets the strength of M cells to enhance the E-M transition rates. The range of θ is set between [1,4] during optimization.

### Growth Competition, Influence (GC_s_I)

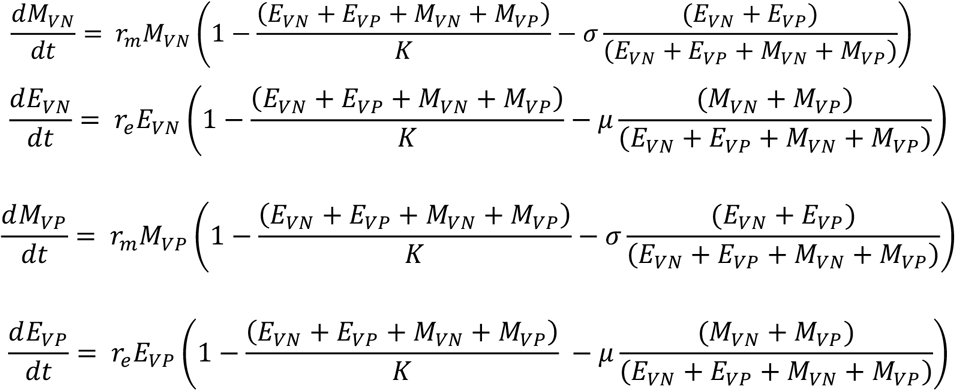

### Growth Competition, Influence and Transition (GC_s_I&T)

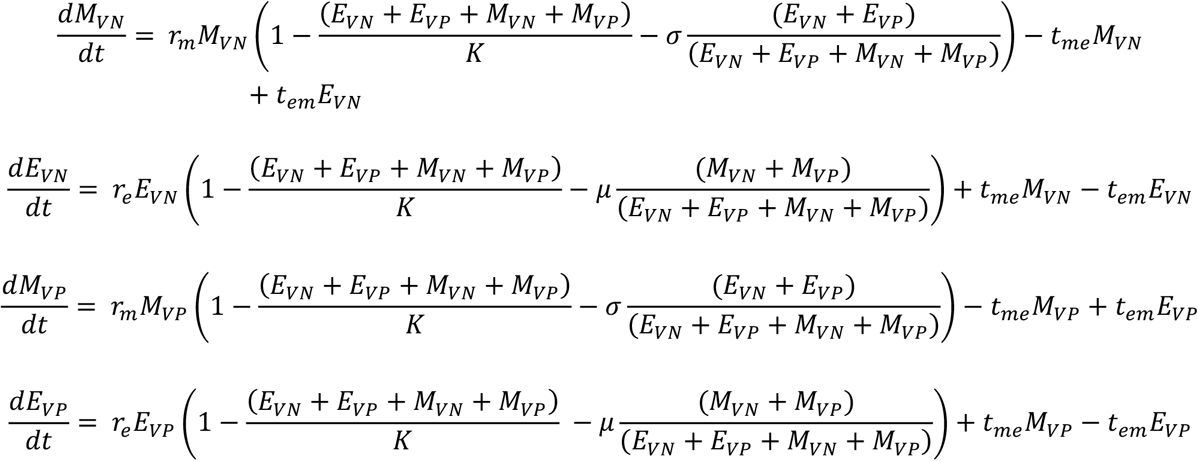

### Parameter identifiability

Parameters can be classified as:

1. Locally Identifiable: the value can be recovered up to finitely many options;
2. Globally Identifiable: the value can be recovered uniquely.

We checked for a priori global identifiability of parameters using a using Differential algebra approach. For the model proposed to fit PMC42-LA experimental data (Table 1) we performed global identifiability calculations analytically (Supplementary File - Identifiability). However, for the more complex models proposed for HCC38 experimental data (Table 2) we used DAISY and StructuralIdentifiability.jl package ^23,25^.

A posteriori local identifiability of model parameter estimates was performed using Profile Likelihood analysis ^24^.

#### Parameter Optimisation

Parameter optimization involves the following steps.

1. Generating trajectories by simulating the population dynamics using the proposed models.
2. Quantifying the difference between the simulated data and the experimental data using a cost/objective (goodness of fit) function. We used the chi-square (χ^2^) values as the cost function:

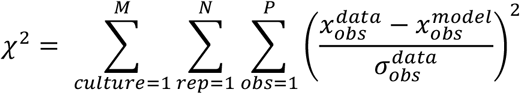 Where, *obs* refer to each experimental observation/measurements, 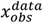, for a given replicate and culture condition with total number of P measurements within the duration of the experiment. The difference between temporal measurements of the experimental data and model simulation is summed for each measurement time point, replicate (rep) and culture condition (culture). 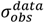 is the standard deviation among the replicates for each measurement time point and the culture condition. M, N and P are number of culture conditions, replicates, and measurement time points, respectively. M = 2, N = 3, and P = 4 for PMC42-LA cells and M = 5, N = 3, and P = 5 for HCC38 cells spontaneous EMP data; and M = 1, N = 3, and P = 5 for HCC38 cells inhibitor treatment data.
3. Searching for the parameter set that minimizes the objective function using an optimizer. The parameter optimization was carried out using the ‘lsqnonlin’ function from MATLAB’s Optimisation toolbox. By default, it uses the ‘trust region reflective’ algorithm, in order to converge to the minimum value of the cost function. The default values of optional parameter of ‘lsqnonlin’ were used for parameter estimation.

### Estimating 95% parameter confidence range and parameter identifiability using Profile Likelihoods

We performed the Profile Likelihood (PL) analysis to estimate the 95% confidence range of optimized model parameters ^24^. The PL analysis uses chi-square statistics to define a threshold level of change in chi-square values from the minimum chi-square values under which model fits are equally good (cost function/goodness of fit). This permissible change in the chi-square from the point of minima depends upon number of model parameters (degree of freedom), and is used to define 95% confidence range for each model parameter. To generate profile likelihood plot of a particular parameter (say θ_5_) in a model of N total parameters (θ_/,…,*N*_),

1. We first choose a relevant range of the parameter (θ_*i*_).
2. Discretize the above parametric range into 1000 values separated by equal steps. Let’s define each value in the discretized interval by 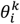, where k = 1 to 1000.
3. Incrementally for each discretized value of θ_*i*_ in its range 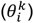, we optimize the other N-1 parameters 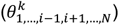 of the model to minimize the chi-square value while keeping the value of 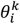 constant.

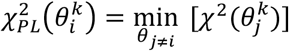
4. The minimum chi-square value thus obtained at each of these points corresponds to the profile likelihood value at the point.
5. While carrying out the optimization for increasing 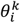 values (k^th^ step), the input guess values for other N-1 parameters 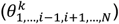 given to the optimization algorithm are taken from the optimized parameter sets for the last step 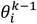 value ((k-1)^th^ step), i.e., 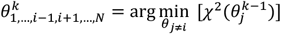

### Generating Synthetic data

To generate synthetic data for a given modal:

1. Multiple parameters set of the model were generated by uniformly sampling each parameter within its lower and upper bounds.
2. For each parameter set, the model dynamics were simulated using the experimental culture condition as the starting point.
3. We then sampled the temporal dynamics of the model output (EpCAM^low^ fraction in models proposed for PMC42-LA cells data, and Venus Cells EpCAM^low^ fraction in models proposed for HCC38 cells data) at limited time points, mimicking the experimental measurement design.
4. Since, for each parameter set for given a model and culture condition we can generate only one temporal trajectory, we added normally distributed noise to the sampled data at each timepoints to get three independent samples, thus mimicking experimental replicates for the culture condition. The standard deviation of normalised distributed noise was adjusted using a denominator term ‘noise factor’. Thus, increasing ‘noise factor’ reduced the variation among replicates of the synthetic data. Further, in cases where adding noise led to negative output (negative E and M fraction) values then resampling of noise term was performed until a positive sum of data with noise term was obtained.
5. We gave same structure to the synthetic data as with the original experimental data for models for HCC38 cells data, i.e., same culture conditions, output measurement timepoints, and number of replicates. For synthetic data generated with models proposed for PMC42-LA cells, the number of measurements timepoints where increased from four to eight, and the culture conditions and replicates were same as the experimental data.

### Improving identifiability of model parameters by adding model informed new data to the existing experimental data

Following are the steps to iteratively identify a timepoint of maximum variability in the output (experimental) variable with respect to parameter of interest and add new (model estimated) data to that time point:

1. For a non-identifiable parameter of a model, we simulate the model dynamics for all parameter sets that are present within the 95% confidence intervals for the non-identifiable parameter, starting with E and M proportions as in the original culture conditions. Note that each point/step in the PL analysis of a parameter corresponds to a set of values of all model parameters. This results in output (Venus cells EpCAM^low^ fraction) variation due to parametric variation. The standard deviation of the output for each culture conditions is calculated at uniformly spaced time points and the standard deviations across culture conditions are then summed up time point wise to give a single time series of standard deviation values for the non-identifiable parameter.
2. The above step is repeated for all non-identifiable parameters of the model to obtain ‘n’ distinct time series of summed standard deviations for the ‘n’ non-identifiable parameters.
3. From the set of ‘n’ distinct time series obtained in step 2, we then find a time point, and the parameter index across ‘n’ time series which has the largest standard deviation values. If the selected time point is already present in the experimental data, then we find another combination of time point and parameter index which results in second (next) largest standard deviation value across ‘n’ time series. The selected time point is the optimal time at which new data will be added using the output variation observed in the selected parameter index.
4. In the output variation observed for each culture condition of the selected parameter index in the step 3, we sample three new output values at the optimal time point. These sampled new values corresponding to the optimal time point for each culture condition are then added to the existing experimental data.
5. With the updated experimental data run the PL analysis for all non-identifiable parameters of step 1 and 2. If any one of the parameters remains non-identifiable then go back to step 1, otherwise stop. The addition of new data is stopped even when a maximum of 4 new data are already added to the original experimental data.

## Author contributions

PJ, RK and MBN performed research, PJ and RK prepared the first draft of manuscript. JTG and MKJ conceived and supervised research and obtained funding. All authors contributed to editing the manuscript and analysed data.

## References

1. Marusyk, A., Janiszewska, M., and Polyak, K. (2020). Intratumor heterogeneity: the Rosetta stone of therapy resistance. Cancer Cell 37, 471. 10.1016/J.CCELL.2020.03.007.

2. Hata, A.N., Niederst, M.J., Archibald, H.L., Gomez-Caraballo, M., Siddiqui, F.M., Mulvey, H.E., Maruvka, Y.E., Ji, F., Bhang, H.E.C., Radhakrishna, V.K., et al. (2016). Tumor cells can follow distinct evolutionary paths to become resistant to epidermal growth factor receptor inhibition. Nature Medicine 2016 22:3 22, 262–269. 10.1038/nm.4040.

3. Pillai, M., Hojel, E., Jolly, M.K., and Goyal, Y. (2023). Unraveling non-genetic heterogeneity in cancer with dynamical models and computational tools. Nature Computational Science 2023 3:4 3, 301–313. 10.1038/s43588-023-00427-0.

4. Lu, M., Jolly, M.K., Levine, H., Onuchic, J.N., and Ben-Jacob, E. (2013). MicroRNA-based regulation of epithelial-hybrid-mesenchymal fate determination. Proc Natl Acad Sci U S A 110, 18144–18149. 10.1073/pnas.1318192110.

5. Hari, K., Ullanat, V., Balasubramanian, A., Gopalan, A., and Jolly, M.K. (2022). Landscape of epithelial mesenchymal plasticity as an emergent property of coordinated teams in regulatory networks. Elife 11. 10.7554/ELIFE.76535.

6. Cook, D.P., and Vanderhyden, B.C. (2020). Context specificity of the EMT transcriptional response. Nat Commun 11, 1–9. 10.1038/s41467-020-16066-2.

7. Chaffer, C.L., Marjanovic, N.D., Lee, T., Bell, G., Kleer, C.G., Reinhardt, F., D’Alessio, A.C., Young, R.A., and Weinberg, R.A. (2013). Poised chromatin at the ZEB1 promoter enables breast cancer cell plasticity and enhances tumorigenicity. Cell 154, 61–74. 10.1016/j.cell.2013.06.005.

8. Panchy, N., Watanabe, K., Takahashi, M., Willems, A., and Hong, T. (2022). Comparative single-cell transcriptomes of dose and time dependent epithelial-mesenchymal spectrums. NAR Genom Bioinform 4, lqac072–lqac072. 10.1093/NARGAB/LQAC072.

9. Yamamoto, M., Sakane, K., Tominaga, K., Gotoh, N., Niwa, T., Kikuchi, Y., Tada, K., Goshima, N., Semba, K., and Inoue, J.I. (2017). Intratumoral bidirectional transitions between epithelial and mesenchymal cells in triple-negative breast cancer. Cancer Sci 108, 1210–1222. 10.1111/cas.13246.

10. Brown, M., Abdollahi, B., and Wilkins, O. (2022). Phenotypic heterogeneity driven by plasticity of the intermediate EMT state governs disease progression and metastasis in breast cancer. Sci Adv 8, eabj8002.

11. Bhatia, S., Monkman, J., Blick, T., Duijf, P.H., Nagaraj, S.H., and Thompson, E.W. (2019). Multi-Omics Characterization of the Spontaneous Mesenchymal–Epithelial Transition in the PMC42 Breast Cancer Cell Lines. J Clin Med 8, 1253. 10.3390/jcm8081253.

12. Bhatia, S., Monkman, J., Blick, T., Pinto, C., Waltham, A., Nagaraj, S.H., and Thompson, E.W. (2019). Interrogation of phenotypic plasticity between epithelial and mesenchymal states in breast cancer. J Clin Med 8, 893.

13. Ruscetti, M., Dadashian, E.L., Guo, W., Quach, B., Mulholland, D.J., Park, J.W., Tran, L.M., Kobayashi, N., Bianchi-Frias, D., Xing, Y., et al. (2016). HDAC inhibition impedes epithelial-mesenchymal plasticity and suppresses metastatic, castration-resistant prostate cancer. Oncogene 35, 3781–3795. 10.1038/onc.2015.444.

14. Biddle, A., Liang, X., Gammon, L., Fazil, B., Harper, L.J., Emich, H., Costea, D.E., and Mackenzie, I.C. (2011). Cancer stem cells in squamous cell carcinoma switch between two distinct phenotypes that are preferentially migratory or proliferative. Cancer Res 71, 5317–5326. 10.1158/0008-5472.CAN-11-1059.

15. Zhou, D., Wu, D., Li, Z., Qian, M., and Zhang, M.Q. (2013). Population dynamics of cancer cells with cell state conversions. Quantitative Biology 1, 201–208. 10.1007/S40484-013-0014-2/METRICS.

16. Wang, Y., Zhou, J.X., Pedrini, E., Rubin, I., Khalil, M., Taramelli, R., Qian, H., and Huang, S. (2023). Cell population growth kinetics in the presence of stochastic heterogeneity of cell phenotype. J Theor Biol 575, 111645. 10.1016/J.JTBI.2023.111645.

17. Vega, S., Morales, A. V., Ocaña, O.H., Valdés, F., Fabregat, I., and Nieto, M.A. (2004). Snail blocks the cell cycle and confers resistance to cell death. Genes Dev 18, 1131–1143. 10.1101/gad.294104.

18. Comaills, V., Kabeche, L., Morris, R., Buisson, R., Yu, M., Madden, M.W., LiCausi, J.A., Boukhali, M., Tajima, K., Pan, S., et al. (2016). Genomic Instability Is Induced by Persistent Proliferation of Cells Undergoing Epithelial-to-Mesenchymal Transition. Cell Rep 17, 2632–2647. 10.1016/j.celrep.2016.11.022.

19. Gollavilli, P.N., Parma, B., Siddiqui, A., Yang, H., Ramesh, V., Napoli, F., Schwab, A., Natesan, R., Mielenz, D., Asangani, I.A., et al. (2021). The role of miR-200b/c in balancing EMT and proliferation revealed by an activity reporter. Oncogene 40, 2309–2322. 10.1038/S41388-021-01708-6.

20. Boareto, M., Jolly, M.K., Goldman, A., Pietilä, M., Mani, S.A., Sengupta, S., Ben-Jacob, E., Levine, H., and Onuchic, J.N. (2016). Notch-Jagged signalling can give rise to clusters of cells exhibiting a hybrid epithelial/mesenchymal phenotype. J R Soc Interface 13. 10.1098/rsif.2015.1106.

21. Gregory, P.A., Bracken, C.P., Smith, E., Bert, A.G., Wright, J. a, Roslan, S., Morris, M., Wyatt, L., Farshid, G., Lim, Y.-Y., et al. (2011). An autocrine TGF-beta/ZEB/miR-200 signaling network regulates establishment and maintenance of epithelial-mesenchymal transition. Mol Biol Cell 22, 1686–1698. 10.1091/mbc.E11-02-0103.

22. Scheel, C., Eaton, E.N., Li, S.H.J., Chaffer, C.L., Reinhardt, F., Kah, K.J., Bell, G., Guo, W., Rubin, J., Richardson, A.L., et al. (2011). Paracrine and autocrine signals induce and maintain mesenchymal and stem cell states in the breast. Cell 145, 926–940. 10.1016/j.cell.2011.04.029.

23. Dong, R., Goodbrake, C., Harrington, H.A., and Pogudin, G. (2022). Differential elimination for dynamical models via projections with applications to structural identifiability.

24. Raue, A., Kreutz, C., Maiwald, T., Bachmann, J., Schilling, M., Klingmüller, U., and Timmer, J. (2009). Structural and practical identifiability analysis of partially observed dynamical models by exploiting the profile likelihood. Bioinformatics 25, 1923–1929. 10.1093/bioinformatics/btp358.

25. Bellu, G., Saccomani, M.P., Audoly, S., and D’Angiò, L. (2007). DAISY: A new software tool to test global identifiability of biological and physiological systems. Comput Methods Programs Biomed 88, 52–61. 10.1016/J.CMPB.2007.07.002.

26. Gupta, P.B., Fillmore, C.M., Jiang, G., Shapira, S.D., Tao, K., Kuperwasser, C., and Lander, E.S. (2011). Stochastic state transitions give rise to phenotypic equilibrium in populations of cancer cells. Cell 146, 633–644. 10.1016/j.cell.2011.07.026.

27. Li, Q., Wennborg, A., Aurell, E., Dekel, E., Zou, J.Z., Xu, Y., Huang, S., and Ernberg, I. (2016). Dynamics inside the cancer cell attractor reveal cell heterogeneity, limits of stability, and escape. Proc Natl Acad Sci U S A 113, 2672–2677. 10.1073/pnas.1519210113.

28. Johnson, K.E., Howard, G.R., Morgan, D., Brenner, E.A., Gardner, A.L., Durrett, R.E., Mo, W., Al’khafaji, A., Sontag, E.D., Jarrett, A.M., et al. (2020). Integrating transcriptomics and bulk time course data into a mathematical framework to describe and predict therapeutic resistance in cancer. Phys Biol 18. 10.1088/1478-3975/abb09c.

29. Pisco, A.O., Brock, A., Zhou, J., Moor, A., Mojtahedi, M., Jackson, D., and Huang, S. (2013). Non-Darwinian dynamics in therapy-induced cancer drug resistance. Nat Commun 4, 1–11. 10.1038/ncomms3467.

30. Shaffer, S.M., Dunagin, M.C., Torborg, S.R., Torre, E.A., Emert, B., Krepler, C., Beqiri, M., Sproesser, K., Brafford, P.A., Xiao, M., et al. (2017). Rare cell variability and drug-induced reprogramming as a mode of cancer drug resistance. Nature 546, 431–435. 10.1038/nature22794.

31. Emert, B.L., Cote, C.J., Torre, E.A., Dardani, I.P., Jiang, C.L., Jain, N., Shaffer, S.M., and Raj, A. (2021). Variability within rare cell states enables multiple paths toward drug resistance. Nature Biotechnology 2021 39:7 39, 865–876. 10.1038/s41587-021-00837-3.

32. Zhou, J.X., Pisco, A.O., Qian, H., and Huang, S. (2014). Nonequilibrium population dynamics of phenotype conversion of cancer cells. PLoS One 9, 1–19. 10.1371/journal.pone.0110714.

33. Risom, T., Langer, E.M., Chapman, M.P., Rantala, J., Fields, A.J., Boniface, C., Alvarez, M.J., Kendsersky, N.D., Pelz, C.R., Johnson-Camacho, K., et al. (2018). Differentiation-state plasticity is a targetable resistance mechanism in basal-like breast cancer. Nat Commun 9. 10.1038/s41467-018-05729-w.

34. West, J., You, L., Zhang, J., Gatenby, R.A., Brown, J.S., Newton, P.K., and Anderson, A.R.A. (2020). Towards multidrug adaptive therapy. Cancer Res 80, 1578–1589. 10.1158/0008-5472.CAN-19-2669/654228/AM/TOWARDS-MULTI-DRUG-ADAPTIVE-THERAPYTOWARDS-MULTI.

35. Marusyk, A., Tabassum, D.P., Altrock, P.M., Almendro, V., Michor, F., and Polyak, K. (2014). Non-cell-autonomous driving of tumour growth supports sub-clonal heterogeneity. Nature 2014 514:7520 514, 54–58. 10.1038/nature13556.

36. Emond, R., Griffiths, J.I., Grolmusz, V.K., Nath, A., Chen, J., Medina, E.F., Sousa, R.S., Synold, T., Adler, F.R., and Bild, A.H. (2023). Cell facilitation promotes growth and survival under drug pressure in breast cancer. Nature Communications 2023 14:1 14, 1–17. 10.1038/s41467-023-39242-6.

37. Pillai, M., Rajaram, G., Thakur, P., Agarwal, N., Muralidharan, S., Ray, A., Barbhaya, D., Somarelli, J.A., and Jolly, M.K. (2022). Mapping phenotypic heterogeneity in melanoma onto the epithelial-hybrid-mesenchymal axis. Front Oncol 12, 913803. 10.3389/FONC.2022.913803/BIBTEX.

38. Paczkowski, M., Kretzschmar, W.W., Markelc, B., Liu, S.K., Kunz-Schughart, L.A., Harris, A.L., Partridge, M., Byrne, H.M., and Kannan, P. (2021). Reciprocal interactions between tumour cell populations enhance growth and reduce radiation sensitivity in prostate cancer. Communications Biology 2021 4:1 4, 1–13. 10.1038/s42003-020-01529-5.

39. Sahoo, S., Mishra, A., Kaur, H., Hari, K., Muralidharan, S., Mandal, S., and Jolly, M.K. (2021). A mechanistic model captures the emergence and implications of non-genetic heterogeneity and reversible drug resistance in ER+ breast cancer cells. NAR Cancer 3. 10.1093/NARCAN/ZCAB027.

40. Sahoo, S., Nayak, S.P., Hari, K., Purkait, P., Mandal, S., Kishore, A., Levine, H., and Jolly, M.K. (2021). Immunosuppressive traits of the hybrid epithelial/mesenchymal phenotype. Front Immunol 12, 797261. 10.3389/fimmu.2021.797261.

41. Harsh Chhajer and Rahul Roy (2023). Rationalised experiment design for parameter estimation with sensitivity clustering. bioRxiv. 10.1101/2023.10.11.561860.

42. Franceschini, G., and Macchietto, S. (2008). Model-based design of experiments for parameter precision: State of the art. Chem Eng Sci 63, 4846–4872. 10.1016/j.ces.2007.11.034.

43. Beik, S.P., Harris, L.A., Kochen, M.A., Sage, J., Quaranta, V., and Lopez, C.F. (2023). Unified tumor growth mechanisms from multimodel inference and dataset integration. PLoS Comput Biol 19, e1011215. 10.1371/JOURNAL.PCBI.1011215.

44. Biddle, A., Gammon, L., Liang, X., Costea, D.E., and Mackenzie, I.C. (2016). Phenotypic Plasticity Determines Cancer Stem Cell Therapeutic Resistance in Oral Squamous Cell Carcinoma. EBioMedicine 4, 138–145. 10.1016/J.EBIOM.2016.01.007.

45. Deshmukh, A.P., Vasaikar, S. V., Tomczak, K., Tripathi, S., Den Hollander, P., Arslan, E., Chakraborty, P., Soundararajan, R., Jolly, M.K., Rai, K., et al. (2021). Identification of EMT signaling cross-talk and gene regulatory networks by single-cell RNA sequencing. Proc Natl Acad Sci U S A 118, e2102050118. 10.73/pnas.2102050118.

46. Pastushenko, I., Brisebarre, A., Sifrim, A., Fioramonti, M., Revenco, T., Boumahdi, S., Keymeulen, A. Van, Brown, D., Moers, V., Lemaire, S., et al. (2018). Identification of the tumour transition states occurring during EMT. Nature 556, 463–468. 10.1038/s41586-018-0040-3.

47. Najafi, A., Jolly, M.K., and George, J.T. (2023). Population dynamics of EMT elucidates the timing and distribution of phenotypic intra-tumoral heterogeneity. iScience 26, 106964. 10.1016/J.ISCI.2023.106964.

48. George, J.T., Jolly, M.K., Xu, S., Somarelli, J.A., and Levine, H. (2017). Survival outcomes in cancer patients predicted by a partial EMT gene expression scoring metric. Cancer Res 77, 6415–6428. 10.1158/0008-5472.CAN-16-3521.

