## Supplementary Information for "Cell-state transitions and frequency-dependent interactions among subpopulations together explain the dynamics of spontaneous epithelial-mesenchymal heterogeneity in breast cancer"

### Analytical Calculation of Identifiability for the models proposed for PMC42-LA experimental data

December 6, 2023

#### Model G

This model is described by the following set of equations:

$$\frac{\mathrm{d}m}{\mathrm{d}t} = r_m m - t_{me} m + t_{em} e - (r_m m + r_e e) m$$

$$\frac{\mathrm{d}e}{\mathrm{d}t} = r_e e + t_{me} m - t_{em} e - (r_m m + r_e e) e$$

Considering

$$\frac{\mathrm{d}m}{\mathrm{d}t} = r_m m - t_{me} m + t_{em} e - (r_m m + r_e e) m$$

$$\Rightarrow \frac{\mathrm{d}m}{\mathrm{d}t} = r_m m - t_{me} m + t_{em}(1-m) - (r_m m + r_e(1-m)) m$$

$$\Rightarrow \frac{\mathrm{d}m}{\mathrm{d}t} = r_m m - t_{me} m + t_{em} - t_{em} m - r_m m^2 - r_e m + r_e m^2$$

$$\Rightarrow \frac{\mathrm{d}m}{\mathrm{d}t} = m^2(r_e - r_m) + m(r_m - t_{em} - r_e - t_{me}) + t_{em}$$

$$\Rightarrow \frac{\mathrm{d}m}{\mathrm{d}t} = am^2 + bm + c$$

By fitting the above reduced model to the data one can estimate parameters a, b, and c. We can then use the values of a, b, and c coefficients to resolve the model parameters. And, by fixing one of the growth rates, we have equal number of equations as unknowns, and thus, we can uniquely determine other growth rate, $t_{me}$, and $t_{em}$ values which makes the model globally identifiable.

#### Model GI

The model is described by the following set of equations:

$$\frac{\mathrm{d}m}{\mathrm{d}t} = r_m(1-\alpha e)m - (r_m(1-\alpha e)m + r_e(1-\beta m)e)m$$

$$\frac{\mathrm{d}e}{\mathrm{d}t} = r_e(1-\beta m)e - (r_m(1-\alpha e)m + r_e(1-\beta m)e)e$$

Considering

$$\frac{\mathrm{d}m}{\mathrm{d}t} = r_m(1-\alpha e)m - (r_m(1-\alpha e)m + r_e(1-\beta m)e)m$$

$$\Rightarrow \frac{\mathrm{d}m}{\mathrm{d}t} = r_m(1-\alpha(1-m))m - (r_m(1-\alpha e)m + r_e(1-\beta m)e)m$$

$$\Rightarrow \frac{\mathrm{d}m}{\mathrm{d}t} = m^3(-\alpha r_m - \beta r_e) + m^2(2\alpha r_m - r_m + \beta r_e + r_e) + m(r_m - \alpha r_m + r_e)$$

$$\Rightarrow \frac{\mathrm{d}m}{\mathrm{d}t} = m^3(-\alpha r_m - \beta r_e) + m^2(-r_m + \beta r_e + r_e) + m(r_m - \alpha r_m + r_e)$$

$$\Rightarrow \frac{\mathrm{d}m}{\mathrm{d}t} = am^3 + bm^2 + cm$$

By fitting the above reduced model to the data one can estimate parameters a, b, and c. We can then use the values of a, b, and c coefficients to resolve the model parameters. And, by fixing one of the growth rates, we have equal number of equations as unknowns, and thus, we can uniquely determine the other growth rate, $\alpha$ and $\beta$ values which makes the model globally identifiable.

### Model G&T-Mr

This model is described by the following set of equations:

$$\frac{\mathrm{d}m}{\mathrm{d}t} = r_m m - t_{me}(1 - \delta m)m + t_{em}e - (r_m m + r_e e)m$$

$$\frac{\mathrm{d}e}{\mathrm{d}t} = r_e e + t_{me}(1 - \delta m)m - t_{em}e - (r_m m + r_e e)e$$

Considering

$$\frac{\mathrm{d}m}{\mathrm{d}t} = r_m m - t_{me}(1 - \delta m)m + t_{em}e - (r_m m + r_e e)m$$

$$\Rightarrow \frac{\mathrm{d}m}{\mathrm{d}t} = r_m m - t_{me}m + \delta t_{em}m^2 + t_{em}(1 - m) - (r_m m + r_e(1 - m))m$$

$$\Rightarrow \frac{\mathrm{d}m}{\mathrm{d}t} = m^2(\delta t_{em} + r_e - r_m) + m(r_m - t_{em} - r_e - t_{me}) + t_{em}$$

$$\Rightarrow \frac{\mathrm{d}m}{\mathrm{d}t} = am^2 + bm + c$$

By fitting the above reduced model to the data one can estimate parameters a, b, and c. We can then use the values of a, b, and c coefficients to resolve the model parameters. And, by fixing $r_m$ and $r_e$, we have equal number of equations as unknowns, and thus, we can uniquely determine $t_{me}$, $t_{em}$ and $\delta$ values which makes the model globally identifiable.

### Model G&T-Mi

This model is described by the following set of equations:

$$\frac{\mathrm{d}m}{\mathrm{d}t} = r_m m - t_{me}m + t_{em}(1 + \theta m)e - (r_m m + r_e e)m$$

$$\frac{\mathrm{d}e}{\mathrm{d}t} = r_e e + t_{me}m - t_{em}(1 + \theta m)e - (r_m m + r_e e)e$$

Considering

$$\frac{\mathrm{d}m}{\mathrm{d}t} = r_m m - t_{me}m + t_{em}(1 + \theta m)e - (r_m m + r_e e)m$$

$$\Rightarrow \frac{\mathrm{d}m}{\mathrm{d}t} = r_m m - t_{me}m + t_{em}(1 + \theta m)(1 - m) - (r_m m + r_e(1 - m))m$$

$$\Rightarrow \frac{\mathrm{d}m}{\mathrm{d}t} = r_m m - t_{me}m + t_{em}(1 - m + \theta m - \theta m^2) - (r_m m^2 + r_e m - r_e m^2)$$

$$\Rightarrow \frac{\mathrm{d}m}{\mathrm{d}t} = m^2(r_e - r_m - \theta t_{em}) + m(r_m + \theta t_{em} - t_{me} - t_{em} - r_e) + t_{em}$$

$$\Rightarrow \frac{\mathrm{d}m}{\mathrm{d}t} = am^2 + bm + c$$

By fitting the above reduced model to the data one can estimate parameters a, b, and c. We can then use the values of a, b, and c coefficients to resolve the model parameters. And, by fixing $r_m$ and $r_e$, we have equal number of equations as unknowns, and thus, we can uniquely determine $t_{me}$, $t_{em}$ and $\theta$ values which makes the model globally identifiable.

## G&T-Er

The model is described by the following set of equations:

$$\frac{\mathrm{d}m}{\mathrm{d}t} = (r_m - t_{me})m + t_{em}(1-\gamma e)e - (r_m m + r_e e)m$$

$$\frac{\mathrm{d}e}{\mathrm{d}t} = (r_e - t_{em}(1-\gamma e))e + t_{me}m - (r_m m + r_e e)e$$

Considering

$$\frac{\mathrm{d}m}{\mathrm{d}t} = (r_m - t_{me})m + t_{em}(1-\gamma e)e - (r_m m + r_e e)m$$

$$\Rightarrow \frac{\mathrm{d}m}{\mathrm{d}t} = (r_m - t_{me})m + (1-\gamma(1-m))(1-m) - (r_m m + r_e(1-m))m$$

$$\Rightarrow \frac{\mathrm{d}m}{\mathrm{d}t} = m^2(-\gamma t_{em} - r_m + r_e) + m(r_m - t_{me} + 2t_{em}\gamma - t_{em} - r_e) + t_{em}(1-\gamma)$$

$$\Rightarrow \frac{\mathrm{d}m}{\mathrm{d}t} = am^2 + bm + c$$

By fitting the above reduced model to the data one can estimate parameters a, b, and c. We can then use the values of a, b, and c coefficients to determine resolve the model parameters. And, by fixing $r_m$ and $r_e$, we have equal number of equations as unknowns, and thus, we can uniquely determine $\gamma$,$t_{em}$ and $t_{me}$ values which makes the model globally identifiable.

#### G&T-EMr

The model is described by the following set of equations:

$$\frac{\mathrm{d}m}{\mathrm{d}t} = (r_m - t_{me}(1-\delta m))m + t_{em}(1-\gamma e)e - (r_m m + r_e e)m$$

$$\frac{\mathrm{d}e}{\mathrm{d}t} = (r_e - t_{em}(1-\gamma e))e + t_{me}(1-\delta m)m - (r_m m + r_e e)e$$

Considering

$$\frac{\mathrm{d}m}{\mathrm{d}t} = (r_m - t_{me}(1-\delta m))m + t_{em}(1-\gamma e)e - (r_m m + r_e e)m$$

$$\Rightarrow \frac{\mathrm{d}m}{\mathrm{d}t} = (r_m - t_{me}(1-\delta m))m + t_{em}(1-\gamma(1-m))(1-m) - (r_m m + r_e(1-m))m$$

$$\Rightarrow \frac{\mathrm{d}m}{\mathrm{d}t} = m^2(t_{me}\delta - t_{em}\gamma + r_m - r_e) + m(r_m - t_{me} - t_{em} + 2\gamma t_{em} + r_e) + t_{em}(1-\gamma)$$

$$\Rightarrow \frac{\mathrm{d}m}{\mathrm{d}t} = am^2 + bm + c$$

By fitting the above reduced model to the data one can estimate parameters a, b, and c. We can then use the values of a, b and c coefficients to resolve the model parameters. But, since even after fixing $r_m$ and $r_e$ we have four unknown variable in three equations, we can not determine $\delta$,$\gamma$,$t_{em}$ and $t_{me}$ values uniquely. Thus this model is not globally identifiable if only $r_m$ and $r_e$ are known.

## G&T-Mi-Er

The model is described by the following equations:

$$\frac{\mathrm{d}m}{\mathrm{d}t} = (r_m - t_{me})m + t_{em}(1 - \gamma e + \theta m)e - (r_m m + r_e e)m$$

$$\frac{\mathrm{d}e}{\mathrm{d}t} = (r_e - t_{em}(1 - \gamma e + \theta m))e + t_{me}m - (r_m m + r_e e)e$$

Considering

$$\frac{\mathrm{d}m}{\mathrm{d}t} = (r_m - t_{me})m + t_{em}(1 - \gamma e + \theta m)e - (r_m m + r_e e)m$$

$$\Rightarrow \frac{\mathrm{d}m}{\mathrm{d}t} = (r_m - t_{me})m + t_{em}(1 - \gamma(1-m) + \theta m)(1-m) - (r_m m + r_e(1-m))m$$

$$\Rightarrow \frac{\mathrm{d}m}{\mathrm{d}t} = m^2(-r_m + r_e - t_{em}(\gamma + \theta)) + m(r_m - t_{me} + t_{em}(2\gamma + \theta - 1) - r_e) + t_{em}(1 - \gamma)$$

$$\Rightarrow \frac{\mathrm{d}m}{\mathrm{d}t} = am^2 + bm + c$$

By fitting the above reduced model to the data one can estimate parameters a, b, and c. We can then use the values of a, b, and c coefficients to resolve the model parameters. But, since even after fixing $r_m$ and $r_e$ we have four unknown variable in three equations, we can not determine $\delta$,$\theta$,$t_{em}$ and $t_{me}$ values uniquely. Thus this model is not globally identifiable if only $r_m$ and $r_e$ are known.

#### Model GI& T

The model is described by the following set of equations:

$$\frac{\mathrm{d}m}{\mathrm{d}t} = r_m(1 - \alpha e)m - t_{me}m + t_{em}e - (r_m(1 - \alpha e)m + r_e(1 - \beta m)e)m$$

$$\frac{\mathrm{d}e}{\mathrm{d}t} = r_e(1 - \beta m)e - t_{em}e + t_{me}m + (r_m(1 - \alpha e)m + r_e(1 - \beta m)e)e$$

Considering

$$\frac{\mathrm{d}m}{\mathrm{d}t} = r_m(1 - \alpha e)m - t_{me}m + t_{em}e - (r_m(1 - \alpha e)m + r_e(1 - \beta m)e)m$$

$$\Rightarrow \frac{\mathrm{d}m}{\mathrm{d}t} = r_m(1 - \alpha(1-m))m - t_{me}m + t_{em}(1-m) - (r_m(1 - \alpha(1-m))m + r_e(1 - \beta m)(1-m))m$$

$$\Rightarrow \frac{\mathrm{d}m}{\mathrm{d}t} = m^3(-\alpha r_m - \beta r_e) + m^2(2\alpha r_m - r_m + r_e + \beta r_e) + m(r_m - \alpha r_m - t_{me} - t_{em} - r_e) + t_{em}$$

$$\Rightarrow \frac{\mathrm{d}m}{\mathrm{d}t} = am^3 + bm^2 + cm + d$$

By fitting the above reduced model to the data one can estimate parameters a, b, c, and d. We can then use the values of a, b, c, and d coefficients to resolve the model parameters. And, by fixing $r_m$ and $r_e$, we have equal number of equations as unknowns, and thus, we can uniquely determine $\alpha$,$\beta$,$t_{em}$ and $t_{me}$ values which makes the model globally identifiable.
