## Supplementary Figures for "Cell-state transitions and frequency-dependent interactions among subpopulations together explain the dynamics of spontaneous epithelial-mesenchymal heterogeneity in breast cancer"

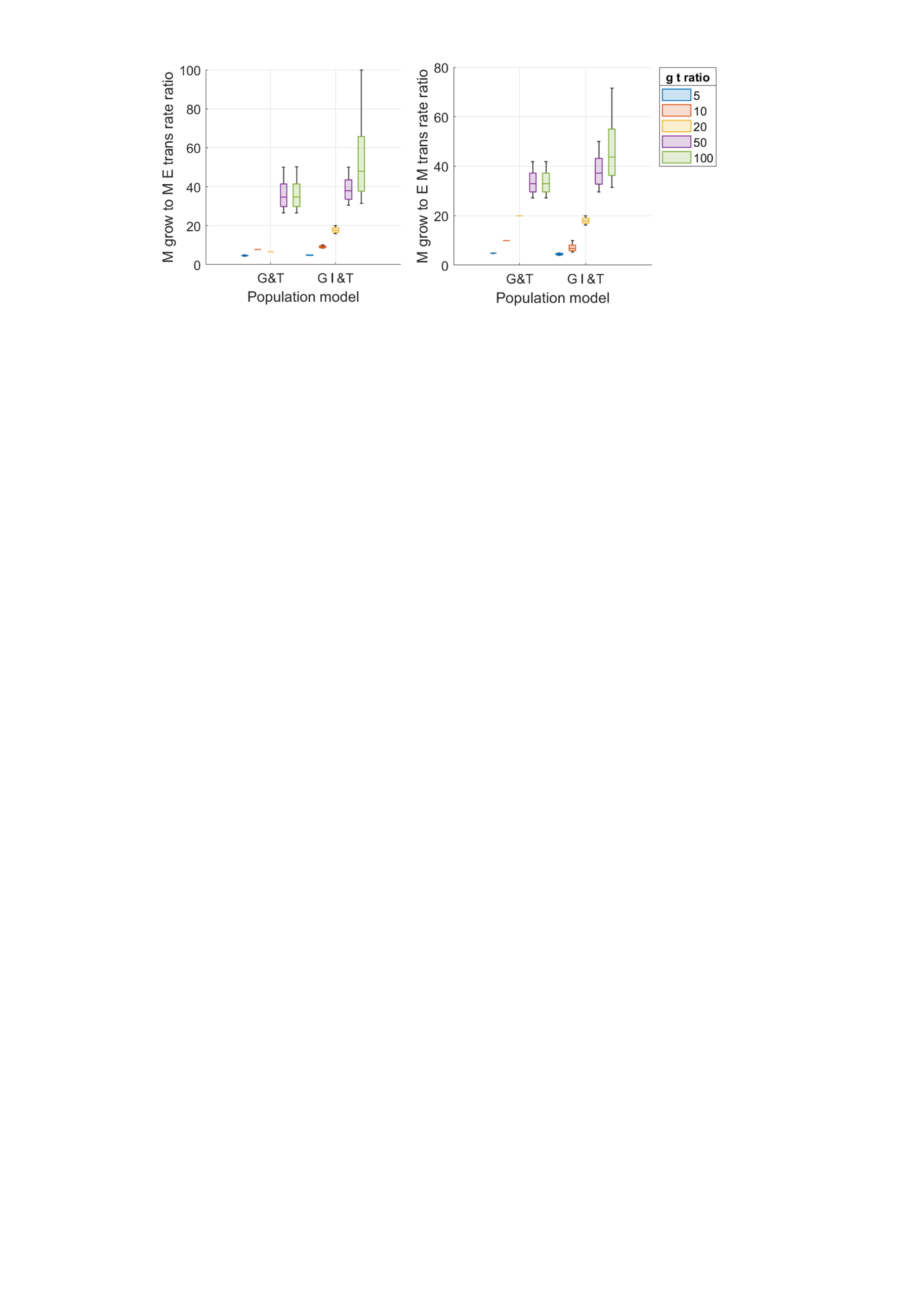

**Figure S1. Ratio of M growth rate to M-E and E-M transition rates for increasing g-t ratio in their 95% confidence range using profile likelihood analysis while fitting G&T and GI&T models to PMC42-LA experimental data**.

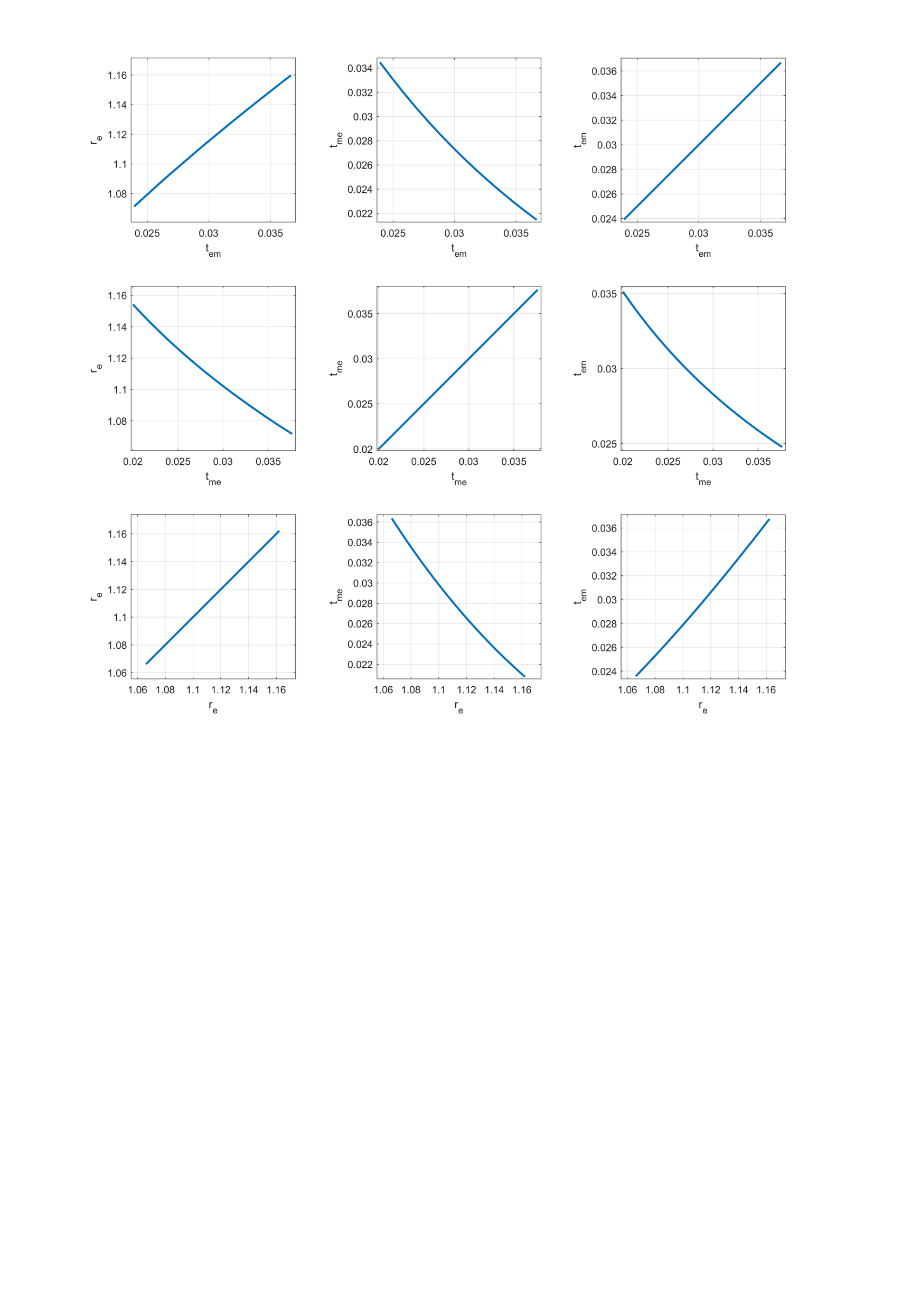

**Figure S2. Parameter co-variability in the 95% confidence range parameter ranges for model G&T while fitting to PMC42-LA experimental data.**

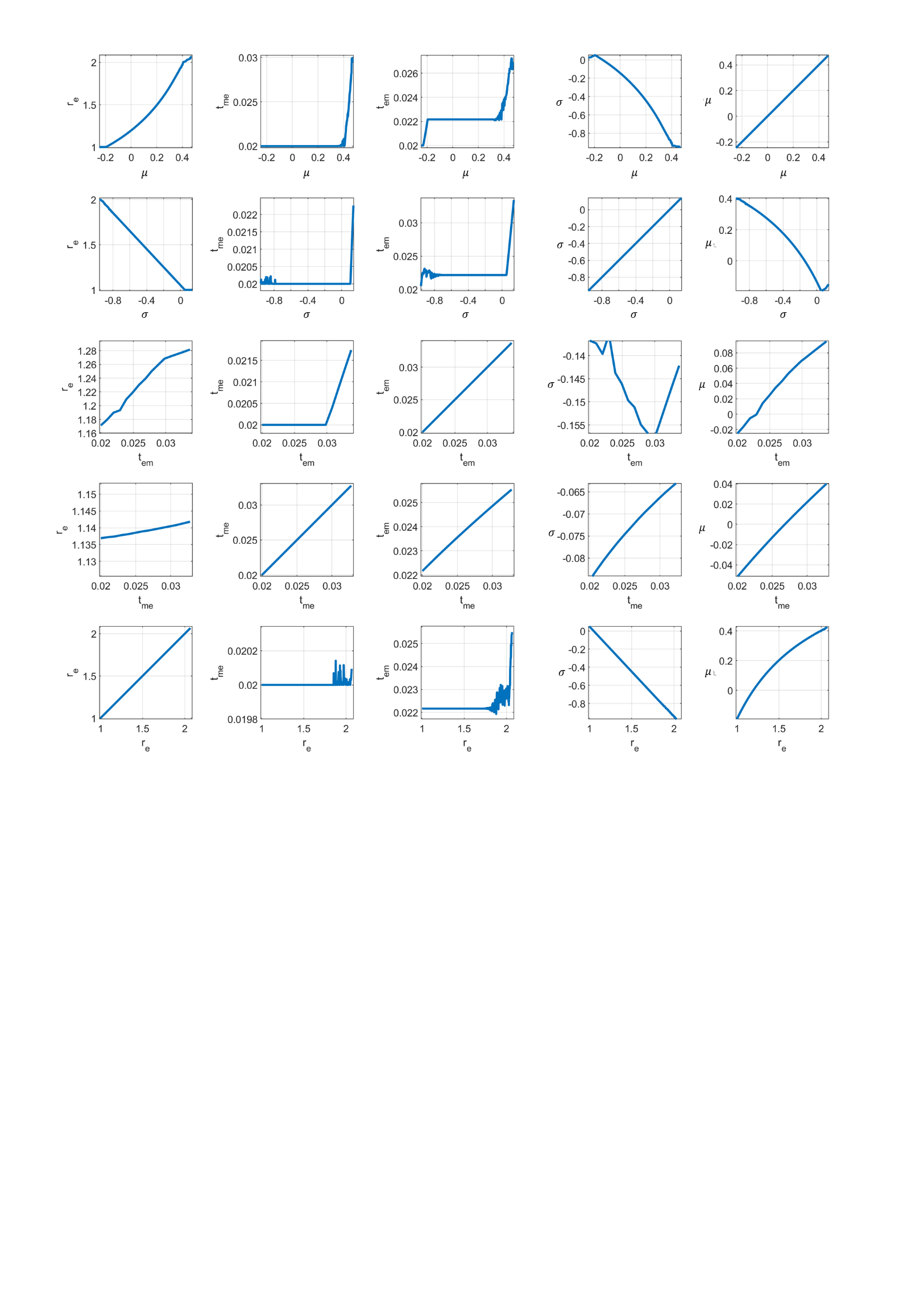

**Figure S3. Parameter co-variability in the 95% confidence range parameter ranges for model GI&T while fitting to PMC42-LA experimental data.**

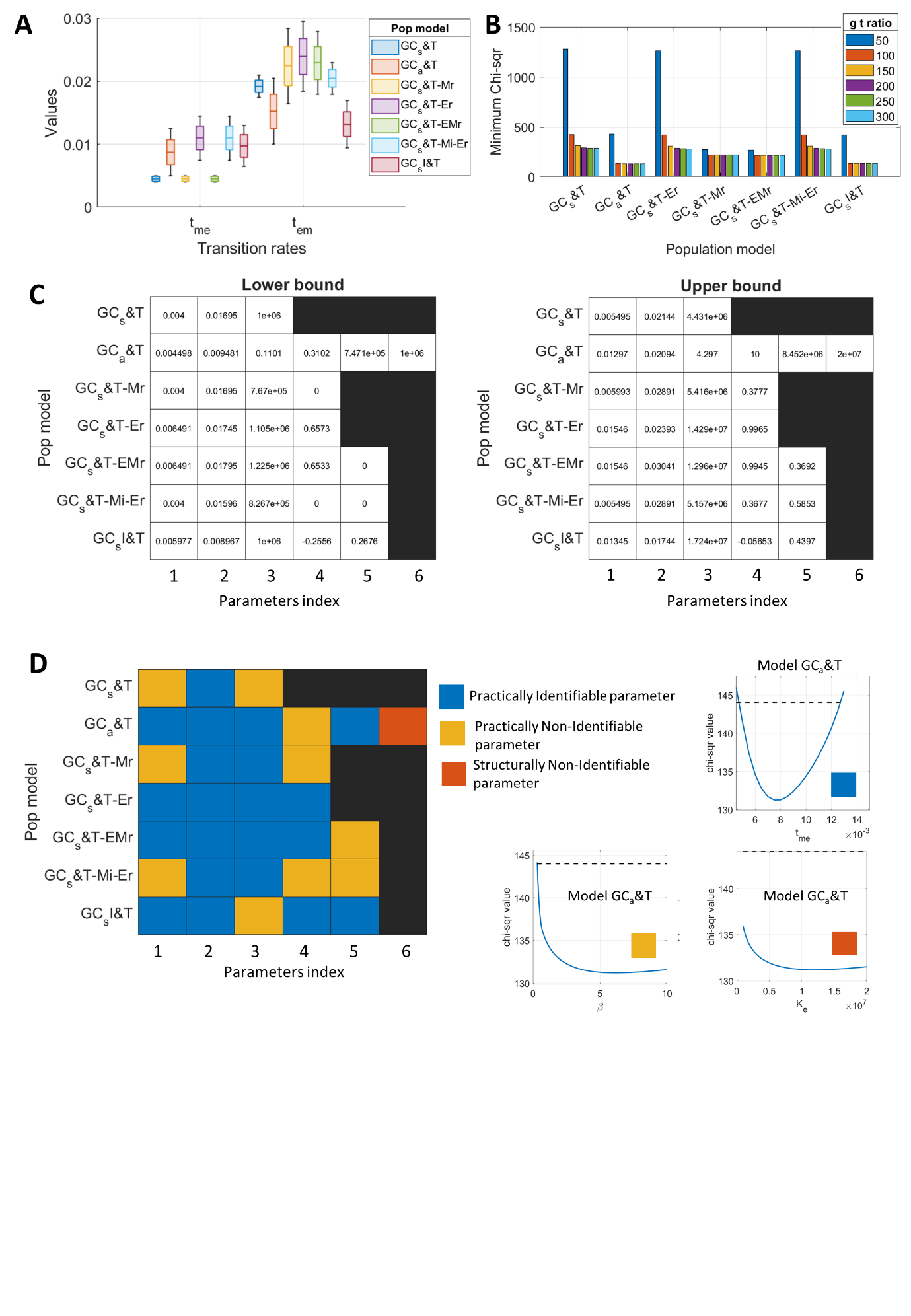

**Figure S4. 95% percent confidence range of parameters for models in Table 2 with increasing g t ratio while fitting the model to HCC38 experimental data (Figure 2A); and parameter identifiability as per the profile likelihood analysis.**  95% percent confidence distribution of **A)** transition rates (t_me_ and t_em_) relative to growth rate of M cells, **B)** Changes in goodness of fit (chi-square) values for increasing g t ratio across population models. **C)** Lower and upper bounds on the estimated parameter as defined by profile likelihood analysis. Refer to Table 6 to map parameter indices on x axis to the parameter variables. **D)** Parameter identifiability for all estimated parameters across models. A parameter’s left/right bound is said to be well defined when the chi-square values cross the threshold (dashed line shown on the right plots) within the search range of the parameter. Otherwise, the parameter has loosely defined left/right bounds. A parameter is practically identifiable when it has both well-defined upper and lower bounds; practically non-identifiable occurs when either left or right bound well defined; and structurally non-identifiable when both left and right bounds are loosely defined.

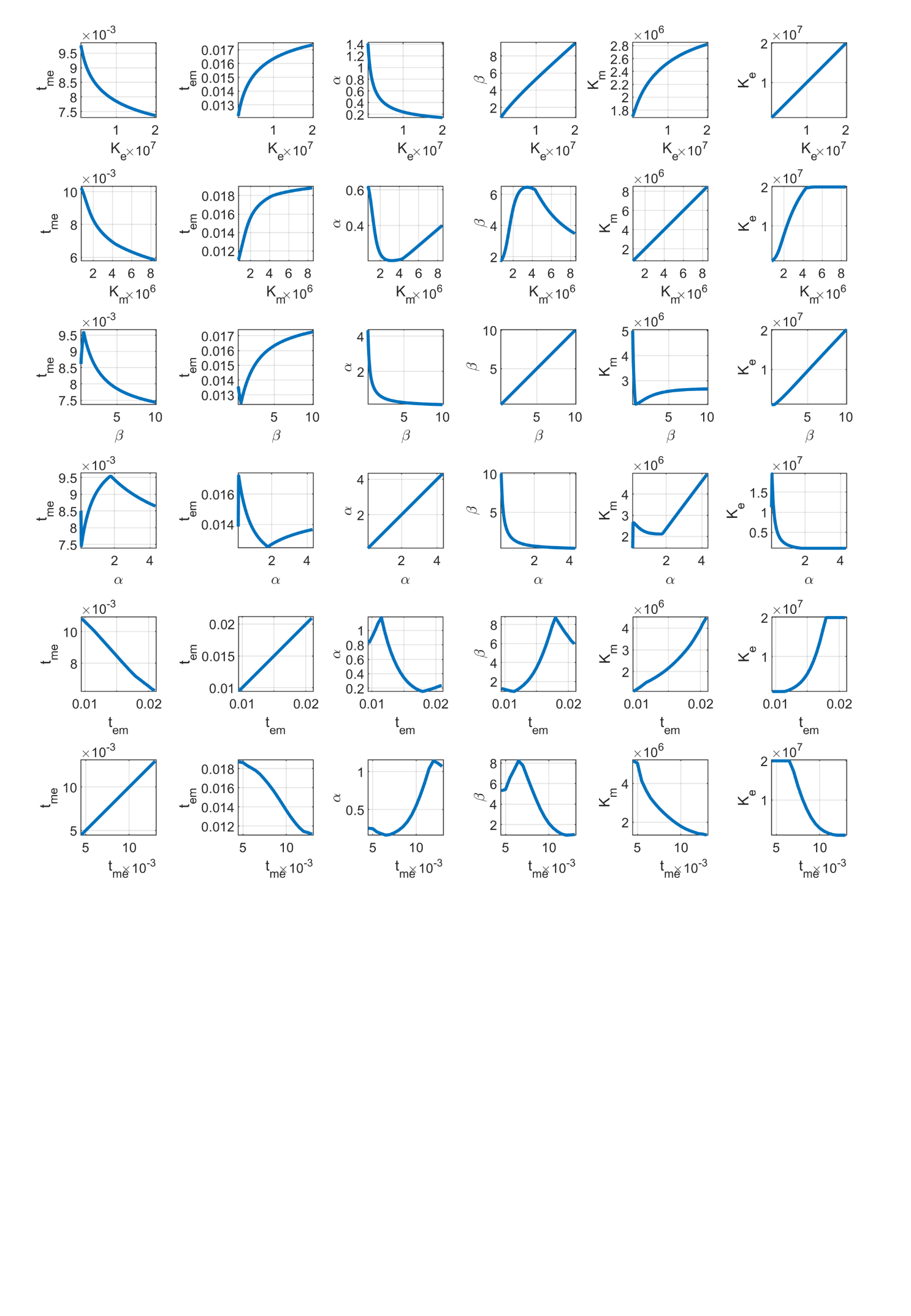

**Figure S5. Parameter co-variability in the 95% confidence range parameter ranges for model GC_a_&T while fitting the model to HCC38 experimental data.**

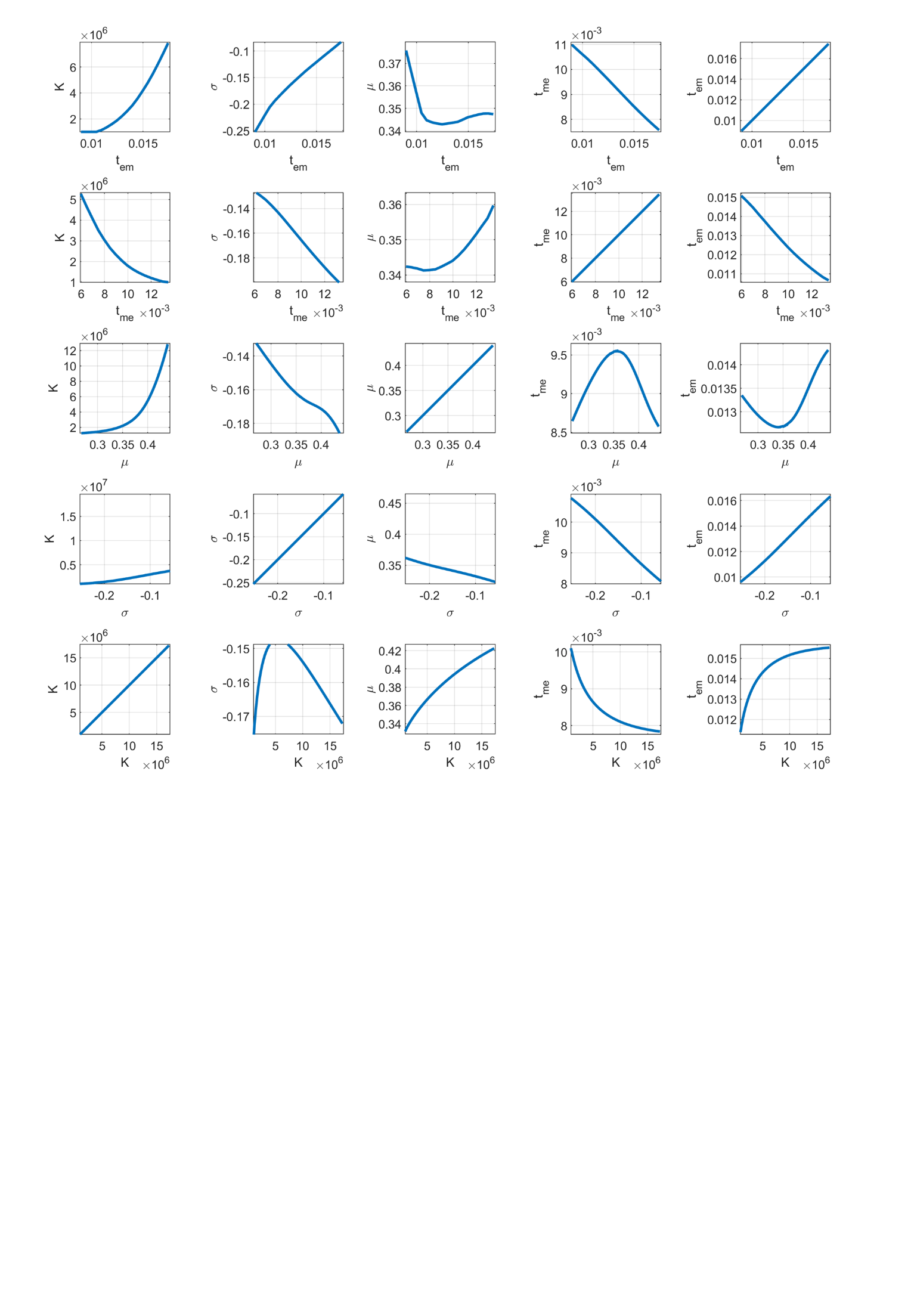

**Figure S6. Parameter co-variability in the 95% confidence range parameter ranges for model GC_s_I&T while fitting the model to HCC38 experimental data.**

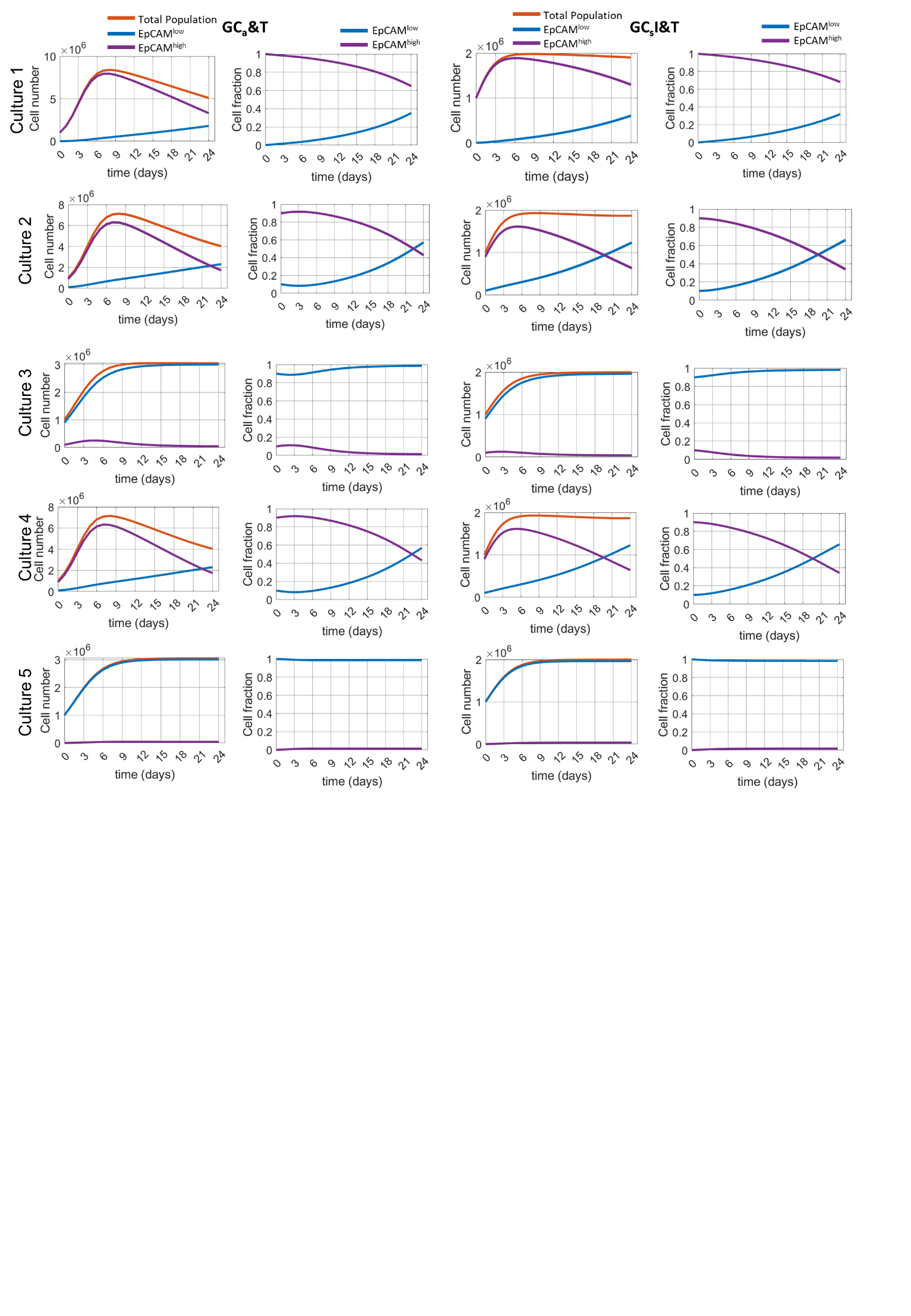

**FigureS7. Temporal cell numbers and population fraction of EpCAM^high^ and EpCAM^low^ cells resulting from GC_a_&T and GC_s_I&T models for best fit parameters.**

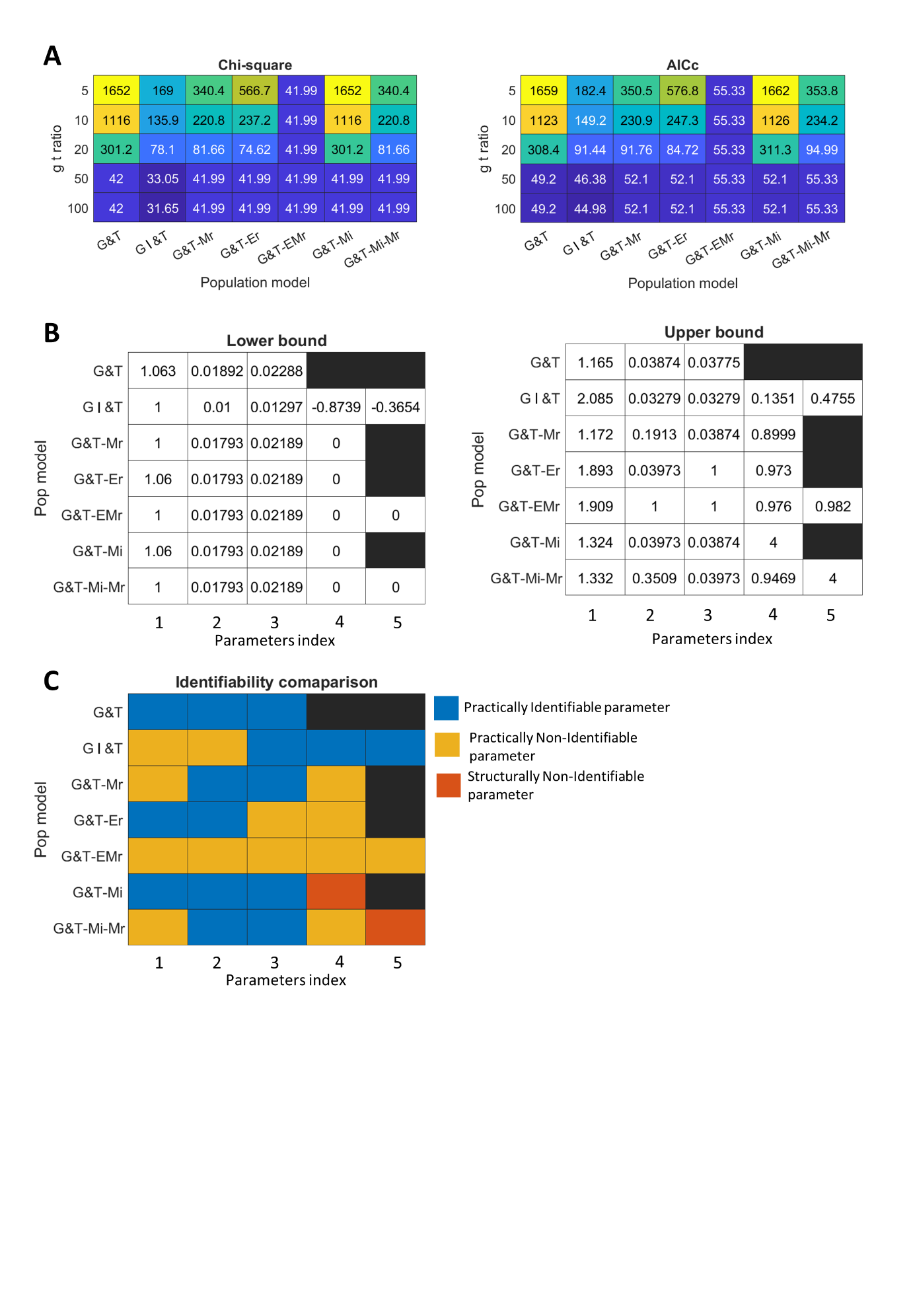

**Figure S8 Goodness of fit and AICc values of models for increasing g-t ratio while fitting the models to PMC42-LA experimental data; and 95% percent bounds on parameters with their identifiability type as per the profile likelihood analysis.** Refer to Table 4 to map parameter indices on x axis in panel B to the parameter variables.

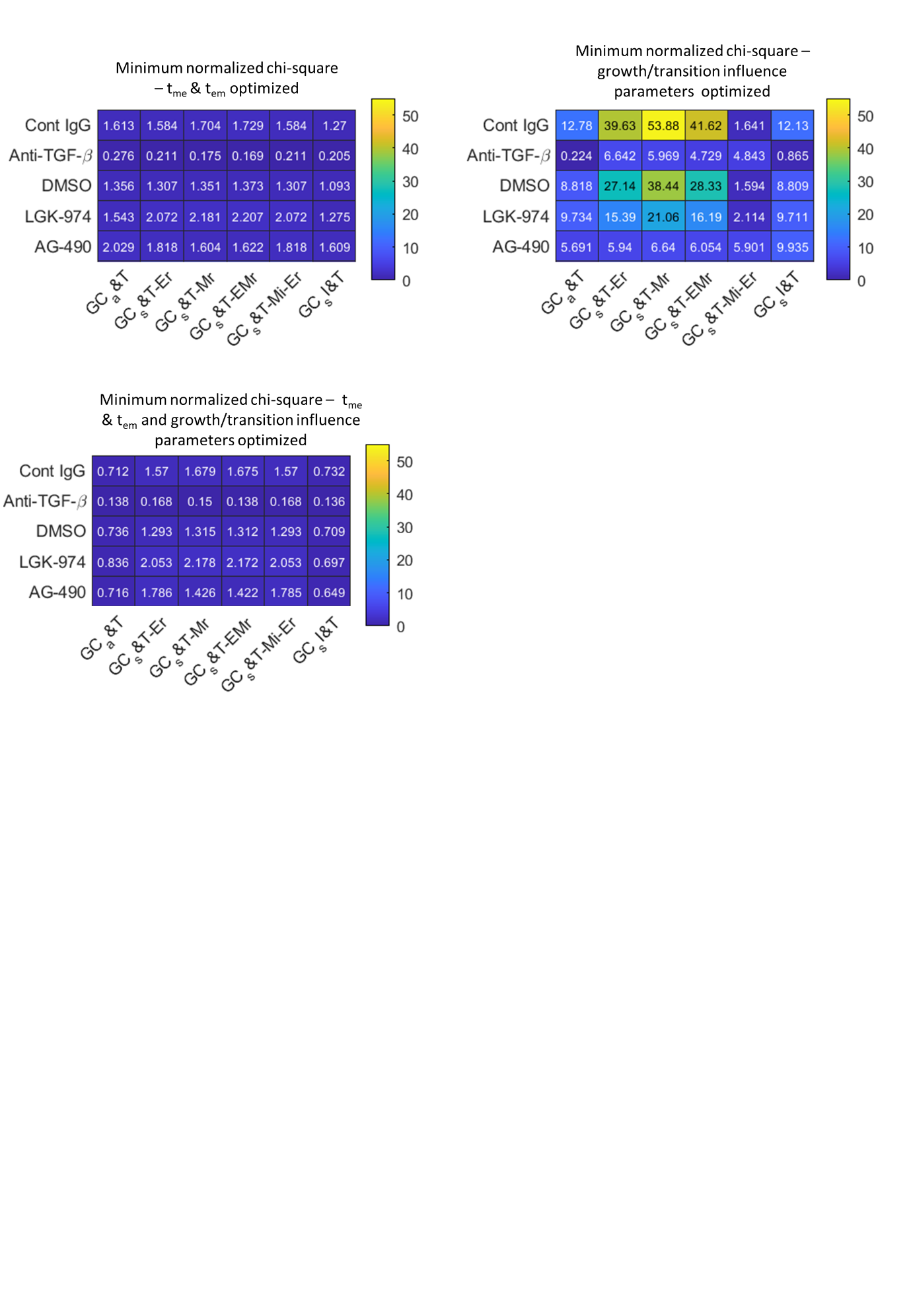

**Figure S9.** Minimum normalized Chi-square values of models proposed for HCC38 experimental data upon fitting to the different inhibitor treatment data.

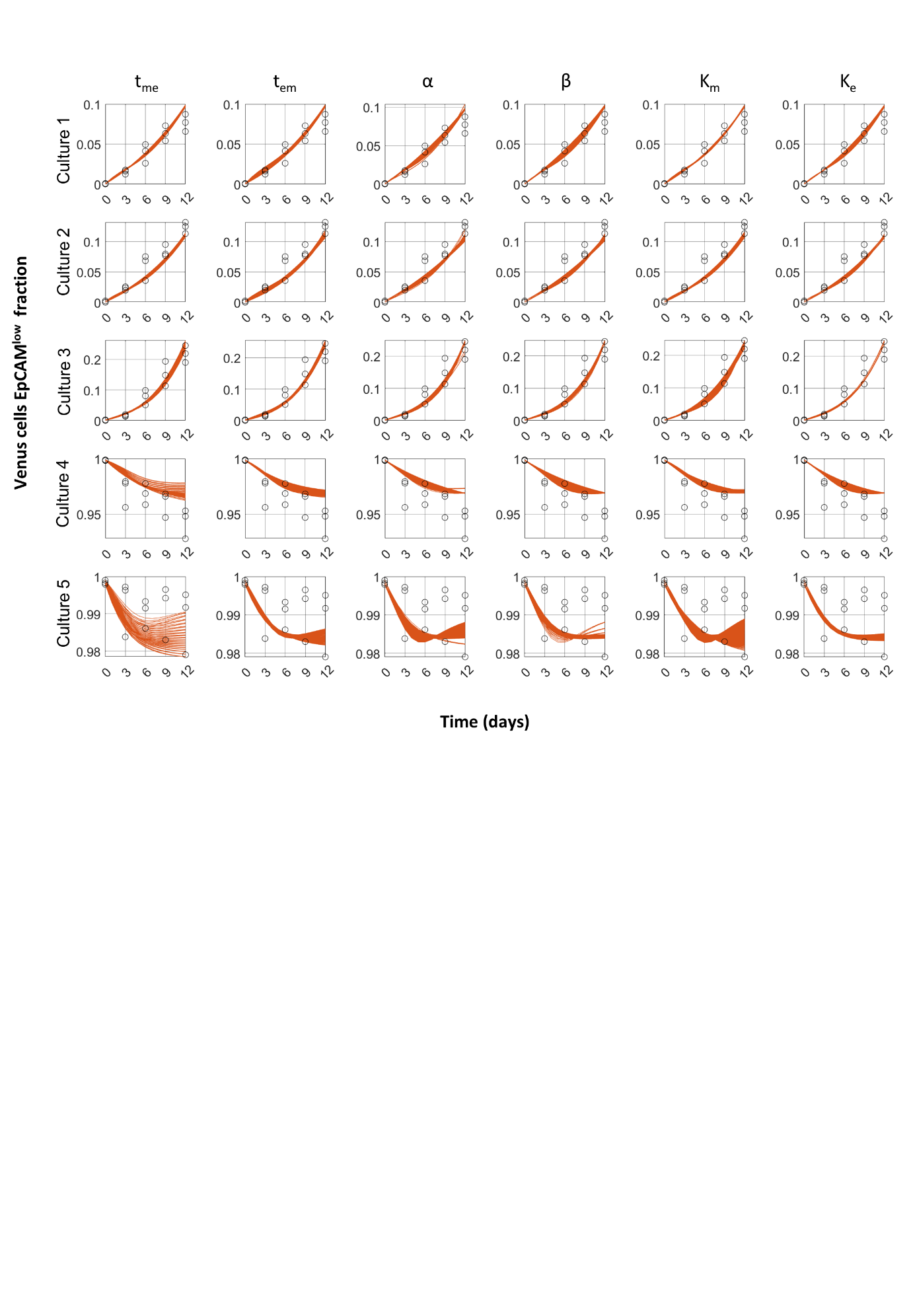

**Figure S10.** Sensitivity of dynamics of Venus EpCAM^low^ cell fraction to variation in GC_a_&T model parameters variation in their 95% confidence range (Figure S4D).
